# HIV-1 promotes cell-to-cell interactions enabling spread from CD4+ T cells to microglia

**DOI:** 10.64898/2026.05.07.723402

**Authors:** F. Reinsberg, K. Schiering, M. G. Lingstaedt, L. Mensching, M. Adiba, T. V. Kraus, J. B. Engler, I. Liebold, L. Bosurgi, S. Schloer, M. Altfeld, M. A. Friese, S. Krasemann, U. C. Lange, W. F. Garcia-Beltran, A. Hoelzemer

## Abstract

HIV-1 infection of the brain occurs early in acute infection and results in neuroinflammation and – when untreated – in cognitive impairment, yet the mechanisms by which microglia become infected remain poorly defined. Evidence from simian immunodeficiency virus (SIV) studies supports a model in which infected CD4+ T cells disseminate HIV-1 to tissue macrophages, but this has not yet been confirmed for human microglia. Here, we used human monocyte-derived microglia (MDMi) and autologous HIV-1-infected primary CD4+ T cells to investigate viral transmission and immune cell interactions. Transcriptional profiling of MDMi confirmed microglia signature genes such as *CX3CR1, P2RY12* and *C1QB*, and surface staining showed expression of CD4 and the HIV-1 coreceptor CCR5. Compared to cell-free infection, direct cell-to-cell contact between MDMi and HIV-1-infected CD4+ T cells markedly enhanced productive infection of MDMi. HIV-1 infection downmodulated the “don’t-eat-me” signal CD47 and increased phosphatidylserine on the surface of primary CD4+ T cells. Consequently, HIV-1 infection of primary CD4+ T cells increased microglia–CD4+ T cell interactions and resulted in enhanced phagocytosis by MDMi. Together, this supports a mechanism where HIV-1 facilitates cell-to-cell spread from primary CD4+ T cells to microglia, which has important implications for therapeutic targeting of HIV-1 brain reservoir seeding.

## Introduction

Untreated human immunodeficiency virus (HIV) infection of the central nervous system (CNS) can result in severe encephalitis due to viral replication in the brain [1, 2]. Even in treated HIV-1 infection, the viral CNS reservoir can drive neuroinflammation causing various degrees of neurocognitive impairment [2–6]. Microglia, the tissue-resident macrophages of the brain, constitute the principal HIV-1 reservoir within the CNS and are a source of productive HIV-1 replication, as recently demonstrated in post-mortem brain samples of people living with HIV-1 [7, 8]. HIV-1 neuroinvasion occurs early after systemic infection [9–11], but the mechanism by which microglia become infected remains unclear. Notably, transmitted/founder HIV-1 variants that dominate during acute HIV-1 infection are predominantly R5 T cell-tropic and require high CD4 surface density on target cells for entry [11, 12], suggesting that microglial infection with cell-free virus may be limited. Due to the difficulty in accessing brain tissue for mechanistic studies in HIV-1 infection, the question of how HIV-1 establishes infection and a reservoir within the brain during acute viremic HIV-1 infection remains unresolved.

Previous studies have shown that circulating CD4+ T cells function as migratory vehicles and disseminate HIV-1 from lymph nodes and peripheral blood to tissue macrophages [13–16]. HIV-1-infected CD4+ T cells engage with myeloid cells in various cell contact-dependent interactions that facilitate viral transmission, overcoming low CD4 surface expression on myeloid cells as a barrier to infection by R5 T cell-tropic HIV-1 variants. First, HIV-1 infection renders CD4+ T cells more susceptible to phagocytosis by macrophages by modulating “eat-me” and “don’t-eat-me” signals, such as surface exposure of phosphatidylserine (PtdSer) [17, 18] and CD47 surface expression [19]. Evidence for infection through phagocytosis – with HIV-infected CD4+ T cells acting as Trojan horses [20] – has been found in both HIV-1 and simian immunodeficiency virus (SIV) models [14–16, 21]. Secondly, heterotypic cell fusion of infected CD4+ T cells and macrophages generates productively infected multinucleated giant cells [22–25]. Thirdly, tunnelling nanotubes (TNT), originally shown to enable HIV-1 transmission between CD4+ T cells over long distances [26], can facilitate HIV-1 spread from CD4+ T cells to macrophages and are implicated in HIV-1 exacerbation in tuberculosis co-infection [27–29].

HIV-1-infected T cells have been proposed as vectors of HIV-1 neuroinvasion as they migrate into the CNS [30–32]. Furthermore, HIV-1-infected T cell clones are shared across the cerebrospinal fluid (CSF) and peripheral blood [33], and pre-treatment with the α4-integrin antagonist Natalizumab, which blocks T-cell entry into the CNS, also blocked brain infection in the SIV rhesus macaque model [34]. Additionally, sequence analyses of HIV-1 RNA isolated from the CSF of people living with HIV showed R5 T cell-tropism during the first months of infection [11] and persistence of HIV-1 RNA in the CSF has been associated with the presence of T cells [35, 36]. Overall, these data from humans and non-human primate studies heavily emphasize a potential role of T cells in mediating HIV-1 infection of the CNS.

As cell-to-cell transmission of HIV-1 is not fully targeted by antiretroviral therapy (ART) and permits replication in virologically suppressed people living with HIV-1 [37, 38], a deeper understanding of whether microglia become infected with HIV-1 via cell-to-cell transfer is critical for guiding the development of therapies that target the CNS reservoir. Here, we established a monocyte-derived microglia model of HIV-1 infection and investigated microglial infection through co-culture with primary HIV-1-infected CD4+ T cells. Our data show that HIV-1 infection enhances the interaction of CD4+ T cells with monocyte-derived microglia, and that direct cell-to-cell contact between these cell types facilitates productive HIV-1 infection of microglia.

## Methods

### PBMC Isolation

Fresh venous blood was collected into EDTA tubes from a donor cohort at the University Medical Center Hamburg-Eppendorf and the Leibniz Institute of Virology after written informed consent. The study was approved by the Ärztekammer Hamburg (PV4780). Peripheral blood mononuclear cells (PBMCs) were isolated from EDTA-anticoagulated whole blood by density gradient centrifugation. Up to 15 mL EDTA blood was transferred to a 50 mL conical tube and diluted 1:1 with HBSS pre-warmed to room temperature; EDTA tubes were rinsed once with HBSS to maximize cell recovery. Alternatively, buffy coat bags were obtained from anonymous healthy adult blood donors through the Mass General Hospital Blood Donor Center (Approval #2022P000699 by the Partners Institutional Review Board/Mass General Brigham Institutional Review Board), poured into a 50 mL conical tube and diluted 1:1 with PBS. The diluted blood was gently layered onto 15 mL LymphoPrep and centrifuged at 950 × *g* for 20 min or 400 × *g* for 30 min (acceleration 3, deceleration 3). The interphase was collected, transferred to a 50 mL conical tube with buffer, and centrifuged at 300 × *g* for 10 min (acceleration 9, deceleration 9) to reduce residual LymphoPrep and platelets. Optionally, the pellet was resuspended in 3 mL ACK lysis buffer for 3 min. Lysis was stopped by adding 2% fetal bovine serum (FBS)/PBS to 40 mL, followed by centrifugation at 500 × *g* for 5 min. Cells were resuspended in RPMI 1640 supplemented with 10% FBS, counted, and cryopreserved at 1–5 × 10⁷ cells/mL.

### Monocyte-derived microglia (MDMi) culture

PBMC-derived monocyte-derived microglia-like cells (MDMi) were generated using a protocol adapted from Quek et al. (2022) [39]. Briefly, on day 0, tissue-culture plates were coated with a thin layer of Geltrex. Geltrex was handled on ice using pre-cooled pipette tips, diluted in ice-cold RPMI-1640 to 100 µg/mL, applied at ∼250 µL/cm², and incubated for 24 h at 37 °C, 5% CO₂. On day 1, MDMi seeding medium (RPMI-1640 containing 10% FBS, 1% GlutaMAX, and 1% penicillin-streptomycin) was prepared and pre-warmed. PBMCs were thawed by adding RPMI-1640 containing 10% FBS (R10) dropwise, counted, and resuspended in seeding medium at ∼1.52 × 10⁶ cells/mL to achieve 4 × 10⁵ cells/cm². After seeding (2.5 mL for 6-well, 0.5 mL for 24-well, 0.3 mL for 48-well plates), plates were gently oscillated and kept at room temperature for 5 min before incubation at 37 °C, 5% CO₂. For differentiation, serum-free RPMI-1640 containing 1% GlutaMAX and 1% penicillin-streptomycin was supplemented fresh with IL-34 (100 ng/mL) and GM-CSF (10 ng/mL) and pre-warmed to 37 °C. On day 2, a full medium change was performed. On days 5, 8, and 11, half-medium changes were performed using a 2× cytokine medium (IL-34 200 ng/mL; GM-CSF 20 ng/mL) by removing half of the culture volume and replacing it with an equal volume of 2× medium. During medium changes, aspirated supernatants were centrifuged and stored at –20 °C for further analysis.

### MDM culture

Monocyte-derived macrophages (MDMs) were generated from PBMCs after positive isolation for CD14+ cells using the STEMCELL CD14+ Isolation Kit. CD14+ cells were thawed, washed, and seeded at 3–5 × 10⁵ cells per well in RPMI-1640 supplemented with 10% FBS, 1% GlutaMAX, and 1% penicillin-streptomycin. CD14+ monocytes were differentiated with either M-CSF (20 ng/mL) as described by Ryan et al. (2017) [40] or GM-CSF (10 ng/mL) following Quek et al. (2022) [41]. Cells were maintained at 37 °C, 5% CO₂ with complete medium changes every 3-4 days. After 7 days, cells were used for downstream applications or maintained in culture with continued complete medium changes every 3-4 days.

### CD4+ T cell culture

CD4+ T cells were negatively isolated from peripheral blood mononuclear cells (PBMCs) using the EasySep™ Human CD4+ T Cell Enrichment Kit (STEMCELL Technologies) or the RosetteSep™ Human T Cell Enrichment Cocktail (STEMCELL Technologies), according to the manufacturer’s instructions. Purified CD4+ T cells were cultured at a density of 1 × 10⁶ cells/mL in RPMI 1640 medium supplemented with 10% FBS, 1% GlutaMAX, 1% penicillin-streptomycin or 1x Primocin, and 100 U/mL interleukin-2 (IL-2) at 37 °C in a humidified atmosphere containing 5% CO₂. After 3-16 h resting, CD4+ T cells were activated by the addition of ImmunoCult™ Human CD3/CD28 T Cell Activator (25 µL/mL; STEMCELL Technologies) or Dynabeads™ Human T-Activator CD3/CD28 for T Cell Expansion and Activation (1 × 10⁶ beads/mL, gibco) to the culture medium. CD4+ T cells were used for downstream analyses or infection experiments 3 days after activation with prior magnetic removal of the Dynabeads where applicable.

### Production of HIV-1 viral stocks

To produce lentiviral particles, HEK293T cells at 70–90% confluence were transfected with the respective full-length HIV-1 containing plasmid using Lipofectamine 3000 transfection reagent (Invitrogen) according to the manufacturer’s protocol. After 48 h, the lentiviral supernatant was collected and centrifuged to remove residual cells and debris. The supernatant was concentrated 100-fold by incubation with either Lenti-X Concentrator (Takara) or PEG-8000 for 1–16 h at 4 °C, and subsequent centrifugation at 1500 × *g* for 45 min to 1 h at 4 °C. The pellet was resuspended in PBS to 1/100 of the original supernatant volume and stored in aliquots at –80 °C until further use. The plasmid containing the proviral genome of the infectious molecular clone (IMC) representing the primary HIV-1 strain CH077 was kindly provided by the Hahn laboratory and the IMC of CH077 was cloned in the Kappes laboratory [42]. The HIV-1 THRO.c-eGFP proviral construct comprising a functional *nef* gene followed by an IRES and the eGFP gene, was kindly provided by the Kirchhoff/Sauter laboratory and generated as previously described [42–44]. The plasmid containing full provirus of an infectious molecular clone of the primary isolate 89.6 was obtained through the NIH HIV Reagent Program [45].

### HIV-1 titration: TZMbl assay

TZM-bl cells were cultured in DMEM (High glucose) supplemented with 10% FBS, 1% Glutamax, 1% penicillin-streptomycin and 1% HEPES at 37 °C and 5% CO₂. For virus titration, 1x 10^4^ TZM-bl cells per well were plated on a 96-well plate and incubated with 40 µg/mL DEAE-Dextran hydrochloride (Sigma-Aldrich) and serial 1:5 dilutions of the concentrated virus stock with a starting dilution of 1:500 for 48 h [46]. BriteLite Plus (PerkinElmer, #6066761) was used as a substrate for a luciferase reporter gene assay and the plate was immediately read out at a TECAN luminometer. TCID50/mL was calculated using the Reed-Muench formula [47]. Plaque-forming units (PFU) per mL were calculated by multiplying TCID50/mL by the conversion factor 0.69 [47].

### Cell-free HIV-1 infection of MDMi

Cell-free HIV-1 infection was conducted by reducing the culture volume of MDMi cultured in six-well plates on day 14 to 1.0 mL (from 2.0 mL), adding 50 µL concentrated HIV-1 stock (multiplicity of infection (MOI) 0.025-0.5), and incubating cells for 6 h at 37 °C, 5% CO₂. Following inoculation, fresh MDMi medium containing 2× cytokines was added at half of the original culture volume. A half-medium change was performed 3 days post-infection. At 6 days post-infection, infection rates were quantified by flow cytometry using antibodies against CD4 and HIV-1 p24.

### HIV-1 infection of CD4+ T cells

Activated CD4+ T cells were resuspended in RPMI 1640 supplemented with 2% FBS at a concentration of 2 × 10⁶ cells/mL. Concentrated HIV-1 was added at a MOI of 0.01–0.1, followed by spinoculation at 1,000 × *g* for 1 h in a 96-well flat-bottom plate. After centrifugation, the medium was replaced with RPMI 1640 supplemented with 10% FBS, 1% GlutaMAX, 1% penicillin-streptomycin, and 100 U/mL IL-2. Cells were cultured at 37 °C in a humidified incubator with 5% CO₂.

### Co-culture of MDMi and HIV-1-infected CD4+ T cells

On day 14 of MDMi culture, 50% of the culture supernatant was removed from each well of the 6-well plates. HIV-1-infected CD4+ T cells (3 days post-infection) were resuspended in MDMi 2x cytokine medium. The CD4+ T cell suspension (3.8 × 10^5^ CD4+ T cells/well) was added to the MDMi cultures either directly or inside a transwell insert (Merck Millipore; pore size 0.4 µm; pore density 100 × 10^6^ pores/cm²). After 24 h of co-culture, the culture medium was aspirated, and wells were washed once with MDMi base medium. Transwell inserts were removed where applicable. Fresh MDMi cytokine medium was then added, and cells were maintained for an additional 6 days at 37 °C and 5% CO₂, with a 50% medium change performed after 3 days. Culture supernatants were collected, centrifuged to remove debris, and stored at −20 °C on days 3 and 6 post co-culture.

### p24 ELISA

Samples from cell culture supernatants were pre-diluted (1:10, 1:100, and 1:1,000) in RPMI 1640. Assays were performed according to the manufacturer’s instructions (Bio-Rad Genscreen™ ULTRA HIV Ag-Ab). Conjugate 1, a biotinylated polyclonal antibody against HIV-1 p24, was added to microplate wells, followed by the addition of pre-diluted samples. After incubation at 37 °C for 1 h and three washes, conjugate 2, streptavidin and peroxidase, was added. Plates were incubated for 30 min at room temperature, washed five times, and developed with substrate solution for 30 min in the dark. The reaction was stopped by adding the stopping solution, and absorbance was measured at 450 nm with a reference wavelength of 620-700 nm. The manufacturer’s HIV-1 p24 antigen standard was used as a reference for quantification.

### Staining and flow cytometry

Adherent cells were gently scraped in 0.5% FBS/PBS, transferred to 96-well plates, and centrifuged at 500 × *g* for 7 min at room temperature between all steps. Myeloid cells were blocked with Fc receptor blocking solution (with or without Live/Dead stain) for 10 min at room temperature in the dark, followed by surface staining in 0.5% FBS/PBS for 30 min at room temperature. Cells were then fixed with 4% paraformaldehyde for 30 min, permeabilized using 1× BD Perm/Wash buffer, and stained with intracellular antibodies for 30 min at room temperature in the dark. After final washes, cells were resuspended in 0.5% FBS/PBS and transferred to cluster tubes. Conventional flow cytometric analysis was performed on a BD FACSymphony A5 analyzer (BD Biosciences), spectral flow cytometric analysis was performed on a Cytek Aurora 5-Laser system (Cytek Biosciences) including unstained controls for baseline fluorescence. Flow cytometry data was analyzed using FlowJo v0.9.0 and v10.10.0 software (Waters Biosciences).

### Annexin V Assay and FACS

CD4+ T cells three days post-infection were washed twice with cold PBS and resuspended in 1× Annexin V Binding Buffer (BD Biosciences). Cells were stained with BV421 Annexin V and 7-AAD (both BD Biosciences) according to the manufacturer’s instructions in the dark at room temperature for 15 minutes. Following incubation, 400 µL of 1× Annexin V Binding Buffer was added to the cell suspension and the samples were analyzed on a BD FACS Fusion cytometer. Annexin V and 7-AAD single and double positive populations were sorted and subsequently stained for intracellular p24 as described above.

### Staining and fluorescence microscopy

Adherent cells were washed with pre-warmed PBS and fixed in 4% paraformaldehyde (PFA) either overnight at 4 °C or for 30 min at 37 °C, followed by three washes in PBS. For overnight in-plate immunofluorescence staining, samples were blocked and permeabilized in 1% BSA and 0.3% Triton X-100 for 45 min at room temperature in the dark. Primary antibodies were diluted in staining buffer containing 0.3% BSA and 0.1% Triton X-100 and incubated with the samples overnight at 4 °C in the dark. The following day, samples were washed three times with PBS, incubated with secondary antibodies (1:500) for 2 h at room temperature in the dark, and counterstained with Hoechst (1:2000 in PBS) for 30 min. After a final wash, samples were mounted with mounting medium, covered with round glass coverslips, and stored at 4 °C protected from light. Images were acquired as z-stacks using a Nikon Ti2 spinning-disk confocal microscope equipped with 405, 488, 561, and 640 nm excitation sources. Images were processed using Fiji software [48].

### Live-cell imaging

For live-cell imaging, MDMi on day 14 of culture (96-well plate) were stained in-plate with CellTrace Far Red. HIV-1 CH077-infected CD4+ T cells (3 days post-infection) were labeled with CellTrace CFSE and were resuspended in RPMI-1640 with a final concentration of 0.5 µM SYTOX Orange. All supernatants were removed from MDMi, and CD4+ T cells were added at 750–24.000 cells/well. The plate was centrifuged for 2 min at 100 × *g* (acceleration 4, deceleration 4) before incubation at 37 °C with 5% CO₂ in the IncuCyte SX5 Live-Cell Analysis System (Sartorius). Images were taken every 10 min for 12 h in the channels Phase, Green, NIR and Orange at 20× magnification.

### MDMi–CD4+ T cell interaction assay

Before the interaction assay, MDMi were washed with MDMi base medium to remove dead cells and debris. Unstimulated (rested in R10 with 100 U/mL IL-2), activated (mock), and HIV-1-infected (89.6, 3 days post-infection) autologous CD4+ T cells were resuspended in MDMi base medium. The respective CD4+ T cells were added to the MDMi at CD4+ T cell-to-MDMi ratios of 2:1 (Donor 1) and 0.5:1 (Donor 2). All wells, including MDMi only controls, were adjusted to a volume of 2 mL with fresh MDMi base medium. The co-culture was incubated for 1 h at 37 °C and 5% CO₂, after which the plates were washed 3 times with PBS to remove unbound CD4+ T cells. For in-plate staining, the cells were blocked with Fc receptor blocking solution for 20 min at 4 °C. Plates were washed 3 times after each blocking, staining or fixation step with 0.5% FBS/PBS or BD Perm/Wash. Surface staining in 0.5% FBS/PBS was added for 30 min at 4 °C in the dark, followed by fixation with BD Cytofix/Cytoperm for 20 min at RT in the dark. Cells were incubated with intracellular staining for 15 min at RT in the dark, washed and stored in 2% FBS/PBS at 4 °C. For both donors, separate plates were prepared for downstream IncuCyte and imaging flow cytometer analysis. A third plate was imaged using the EVOS M5000 Imaging System (Invitrogen), then cells were detached by gentle scraping and analyzed by a conventional cytometer.

### IncuCyte analysis

For image acquisition of stained (in-plate) and fixed cells, an IncuCyte SX5 (Sartorius) was used. 49 images per well were taken in the channels Phase, Green, NIR and Orange at 20× magnification. Segmentation was performed using phase images and MDMi were selected based on area and eccentricity parameters. Subsequent gating of cells was performed based on the mean fluorescence intensity of the markers CD3 and p24.

### Imaging flow cytometry (ImageStream)

Stained cells were acquired on a Cytek Amnis ImageStream^X^ Mk II Imaging Flow Cytometer (Amnis, Cytek Biosciences) with a 60× objective. Fluorescence compensation was performed using FlowJo software. Subsequent image analysis and visualization were done with IDEAS 6.2 software (Amnis, Cytek Biosciences).

### MDM–CD4+ T cell interaction assay

MDM generated with M-CSF were washed with R10 to remove dead cells and debris. Autologous unstimulated CD4+ T cells were thawed the day before and incubated overnight in R10 with 50 U/mL IL-2. For antibody coating, a fraction of the unstimulated CD4+ T cells was incubated with a CD4 antibody (10 µg/mL) for 15 min at 37 °C and then washed. Unstimulated, antibody-coated, and activated CD4+ T cells were resuspended in R10. The respective CD4+ T cells (or R10 for MDMi only controls) were added to the MDM in a 2:1 ratio of CD4+ T cells to MDM. The co-culture was incubated for 1 h at either 37 °C and 5% CO₂ or at 4 °C. To get rid of unbound CD4+ T cells, the plates were washed with PBS. For in-plate staining, the cells were incubated with Fc receptor blocking/LD staining mix for 30 min at 4 °C. After each blocking, staining or fixation step, plates were washed 3 times with 0.5% FBS/PBS or BD Perm/Wash. Cells were stained with surface staining in 0.5% FBS/PBS for 30 min at 4 °C in the dark, followed by fixation with BD Cytofix/Cytoperm for 20 min at RT in the dark. Subsequently, cells were stained with intracellular staining for 15 min at RT in the dark, washed and stored in 2% FBS/PBS at 4 °C. For flow cytometry analysis, cells were gently scraped.

### RNA isolation

For RNA isolation, MDMi were cultured in 6-well plates and RNA was extracted using the RNeasy Micro Kit (Qiagen). Briefly, day 14 MDMi and day 7 M-CSF-derived MDM were washed with PBS and lysed in 250 µL RLT buffer supplemented with 1% β-mercaptoethanol. Adherent cells were scraped and transferred to 1.5 mL microcentrifuge tubes. For suspension samples, 2 × 10⁶ cells were resuspended directly in RLT buffer supplemented with 1% β-mercaptoethanol. Lysates were stored at −80 °C until extraction. For extraction, lysates were adjusted to ≤5 × 10⁵ cells in 350 µL RLT buffer. For positively isolated samples, magnetic beads were removed using a DynaMag-2 before column loading. Lysates were mixed 1:1 with 70% ethanol and processed on RNeasy MinElute spin columns with on-column DNase digestion (15 min) according to the manufacturer’s instructions, including washes with RW1 and RPE and a final 80% ethanol wash. Columns were air-dried and RNA was eluted in 20 µL RNase-free water, quantified by NanoDrop, and stored at −80 °C.

### Bulk RNA sequencing and data analysis

Total RNA was isolated from monocytes, M-CSF-derived macrophages (MDM), and MDMi from three donors. RNA quality was assessed using an Agilent Bioanalyzer (RNA 6000 Nano). For each sample, 300 ng total RNA was subjected to poly(A) selection (NEBNext Poly(A) mRNA Magnetic Isolation Module), followed by library preparation (Lexogen CORALL Total RNA-Seq Library Prep with UDI Set B1) according to the manufacturers’ protocols. Libraries were quality controlled and sequenced on an Illumina NextSeq 2000 (P3, 100-cycle kit) in paired-end mode (2 × 57 bp). Reads were demultiplexed with bcl2fastq, yielding 68.3–82.6 million read pairs per sample, and assessed with FastQC v0.12.0. Reads were aligned to the human reference genome GRCh38 (Ensembl release 108) using STAR v2.7.10b, converted to BAM with samtools, and gene-level counts were generated with HTSeq. RNA-seq analysis. Downstream analyses were performed in R (v4.4.2) using DESeq2; PCA, volcano plots, and heatmaps were generated with ggplot2, ggrepel, and pheatmap. For comparative PCA, in-house samples and published reference samples from Abud et al. (2017) [49] were analyzed jointly after variance-stabilizing transformation, and the first two principal components were calculated from the 500 most variable genes. For the donor-matched dataset, differential expression between MDMi and MDM was tested with a donor-blocked design using Wald tests with Benjamini-Hochberg correction (adjusted P < 0.05, |log₂ fold change| ≥ 1). Shrunken fold-change estimates (ashr) were used for volcano plots. Heatmaps show row-scaled variance-stabilized expression of selected immune and phagocytosis receptor genes, with samples ordered by cell type and donor and genes hierarchically clustered. Gene identifiers were mapped using AnnotationDbi and org.Hs.eg.db.

### Statistical analysis

GraphPad Prism 10.61 (GraphPad Software, Inc.) was used for data visualization and statistical analyses. Statistical comparisons were performed using the Wilcoxon matched-pairs signed-rank test. Correlation was assessed using a two-tailed Spearman’s rank correlation coefficient. P values < 0.05 were considered statistically significant.

## Results

### Monocyte-derived microglia-like cells exhibit homeostatic microglial features and are susceptible to HIV-1 infection

Given the limited availability of primary human microglia for mechanistic studies of HIV-1 infection, we investigated a peripheral blood monocyte-derived human microglia-like model as a system for HIV-1 infection. The monocyte-derived microglia-like cell (MDMi) model published by Quek et al. (STAR Protocols 2022) robustly generated a ramified microglia-like phenotype from peripheral blood mononuclear cells (Figure 1A) that could be cultured up to 21 days. MDMi expressed the leukocyte and myeloid markers CD45, CD11b (Supplemental Figure S1A, B), CD68 and Iba-1 (Figure 1A). Notably, P2RY12, marking homeostatic microglia, was selectively expressed in MDMi, and clearly distinguished these cells from the monocyte-derived macrophage (MDM) control conditions differentiated with either GM-CSF or M-CSF (Figure 1A). Stimulation of MDMi with either lipopolysaccharide (LPS)/ Interferon-γ or adenosine triphosphate (ATP) confirmed functional downregulation of the P2RY12 protein on MDMi upon activation (Supplemental Figure S1C). Compared to M-CSF-differentiated but not to GM-CSF-differentiated MDM, MDMi expressed similar surface levels of the chemokine receptor CX3CR1 that interacts with neuron-derived fractalkine (Figure 1A). To further assess the transcriptomic profile of MDMi, we performed bulk RNA-seq on MDMi, M-CSF-derived MDMs, and monocytes from peripheral blood, and compared these data with a published dataset from Abud et al. (2017) [49], which includes peripheral myeloid cells, iPSC-derived microglia, and primary adult and fetal human microglia. Principal component analysis of the top 500 variable genes clearly separated monocytes from the microglial reference samples and positioned MDMi next to primary adult and fetal microglia (Figure 1B). However, MDMi clustered most closely with MDMs and remained distinct from both primary microglial populations, indicating a partial rather than complete acquisition of a microglia-like transcriptional identity. For example, both the monocyte-derived microglia model and the iPSC-derived model lacked *TMEM119* expression, a marker of resting microglia in the human brain. Notably, the iPSC-derived microglia in the published dataset from Abud et al. formed a discrete cluster, separate from both primary microglia and MDMi from this study. Despite their proximity to MDMs in PCA space, differential expression analysis revealed selective transcriptional differences between MDMi and MDMs, with significantly higher expression of several human microglia-associated signature genes in MDMi [50], including *P2RY12, P2RY13, CX3CR1, C1QB,* and *PROS1* (Supplemental Figure S1D).

**Figure 1:**
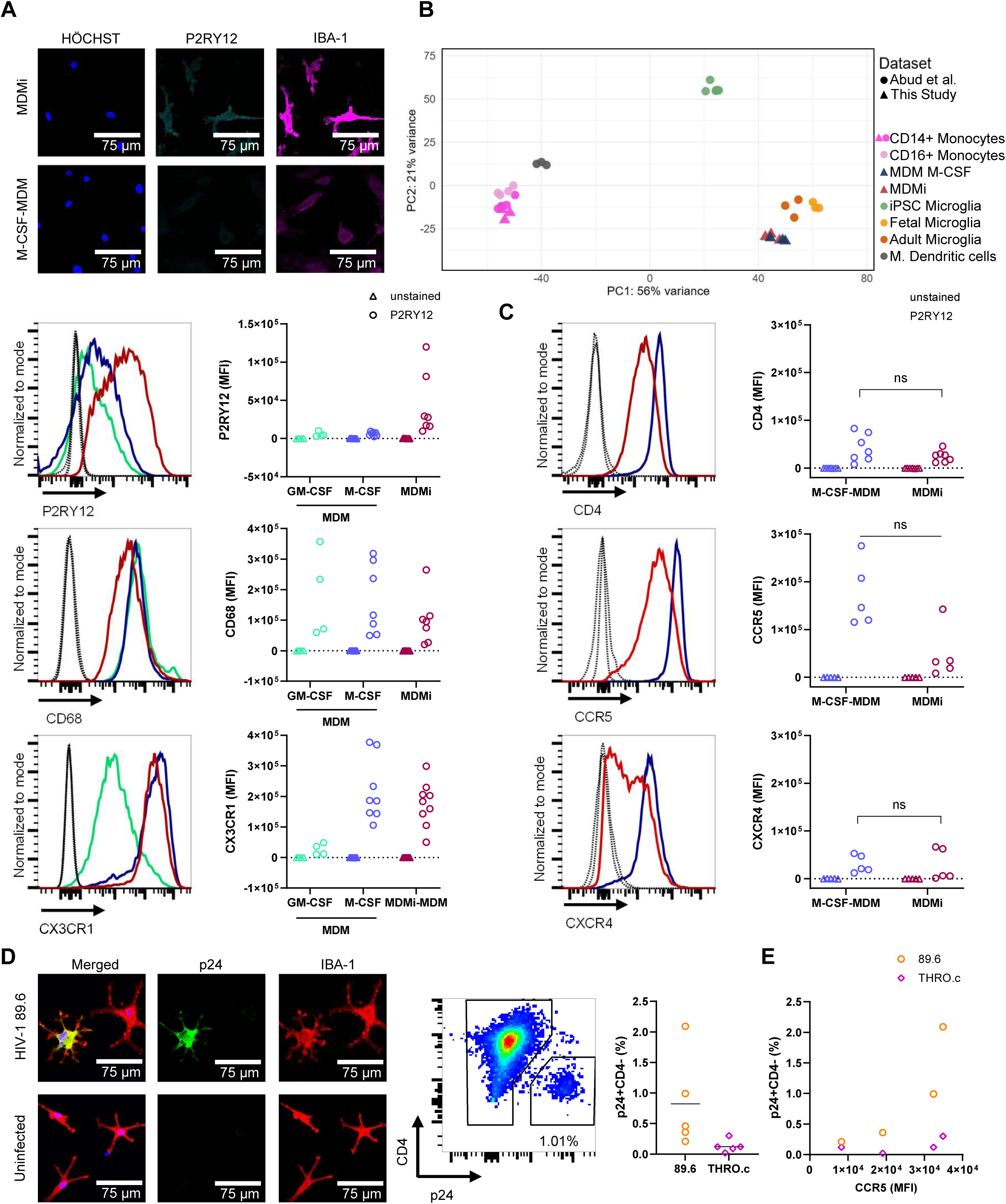
Monocyte-derived microglia exhibit homeostatic microglial features and are susceptible to cell-free HIV-1 infection. (A) Upper panel: representative immunofluorescence images of monocyte-derived microglia-like cells (MDMi) and monocyte-derived macrophages differentiated with M-CSF (MDM) stained for Hoechst, P2RY12, and IBA-1. Lower panel: representative flow cytometric histograms (dotted lines = unstained controls) and median fluorescence intensity (MFI) of P2RY12, CD68, and CX3CR1 expression in GM-CSF-derived MDM, M-CSF-derived MDM, and MDMi (triangles = unstained controls; circles = stained samples). (B) Principal component analysis of bulk RNA-seq profiles from monocytes, M-CSF-derived MDM, and MDMi (top 500 variable genes; n = 3 donors; triangle symbol) compared with peripheral myeloid and microglial reference populations published by Abud et al. (circle symbol) [49]. (C) Representative flow cytometric histograms (dotted lines = unstained controls) and cumulative data (MFI) of surface CD4, CCR5, and CXCR4 expression on M-CSF-derived MDM and MDMi (triangles = unstained controls; circles = stained samples). Statistical significance was assessed using the Wilcoxon matched-pairs signed-rank test (ns – not significant). (D) Representative immunofluorescence images of 89.6-infected and uninfected MDMi stained for HIV-1 p24, IBA-1, and Hoechst (scale bars, 75 µm), representative flow cytometric gating of infected cells based on p24 positivity and CD4 downregulation, and quantification of productive infection as the percentage of p24+CD4− cells after infection with dual-tropic 89.6 or the R5-tropic reporter virus THRO.c.GFP (n = 5 donors). (E) CCR5 surface expression (MFI) and infection frequency in MDMi (n = 4 donors). Correlation was assessed using a two-tailed Spearman’s rank correlation coefficient (not significant).

As a next step, we investigated the expression of CD4 as a HIV-1 entry receptor and CCR5 and CXCR4 as HIV-1-coreceptors on the surface of the MDMi model (Figure 1C). MDMi expressed CCR5, as well as low levels of CXCR4. CD4 and CCR5 levels did not differ significantly between MDMi and MDM. Accordingly, we next assessed cell-free infection using an R5-tropic transmitted founder virus (THRO.c.) [42] and the dual-tropic HIV-1 strain 89.6 that was isolated from an individual with advanced HIV-1 disease [45]. Both viruses have been shown to infect monocyte-derived macrophages [42, 44]. We observed productive HIV-1 infection of MDMi by both strains, as assessed via p24 capsid staining and functional CD4 downregulation on infected MDMi 6 days post infection (Figure 1D). Infection frequency, measured as percentage of p24+CD4-cells, was low and ranged from 0.21 to 2.09% for 89.6 and from 0.02 to 0.3% for THRO.c. Higher surface expression of the HIV-1 coreceptor CCR5 on MDMi was associated with a higher infection frequency (Figure 1E), although this association was not statistically significant. Taken together, the MDMi model expressed microglia signature markers such as P2RY12 and CX3CR1, showed transcriptional similarity to primary microglia and M-CSF-derived MDMs, and was susceptible to cell-free HIV-1 infection.

### Cell contact between HIV-1-infected CD4+ T cells and MDMi enhances productive MDMi infection

As infection of myeloid cells with T-tropic HIV-1 variants – that dominate during acute infection [11] – is limited by the low CD4 density on myeloid cells [12], we tested whether cell-to-cell transmission of HIV-1 from infected CD4+ T cells facilitates microglia infection. To differentiate cell-free infection from cell contact-dependent transmission, MDMi were either directly co-cultured with autologous HIV-1-infected CD4+ T cells or separated by a transwell insert with a pore size of 0.4 µm that allowed for HIV-1-virion but not immune-cell migration (Figure 2A). CD4+ T cells were infected with the dual-tropic HIV-1 variant 89.6 and the transmitted founder strain CH077 that is R5 and weakly X4-tropic [42] to allow for cell-free virus infection. Infection frequencies of primary CD4+ T cells ranged from 9.8 to 26.9% for 89.6 and 1.26 to 11.0% for CH077 three days post-infection before MDMi co-culture (Figure 2B). After 24-hour co-culture with MDMi, HIV-1-infected CD4+ T cells were removed from the culture plates and the MDMi culture was continued for six additional days to assess productive infection of MDMi via flow cytometry by functional downregulation of CD4 in HIV-1 p24+ cells (Figure 2A, C).

**Figure 2:**
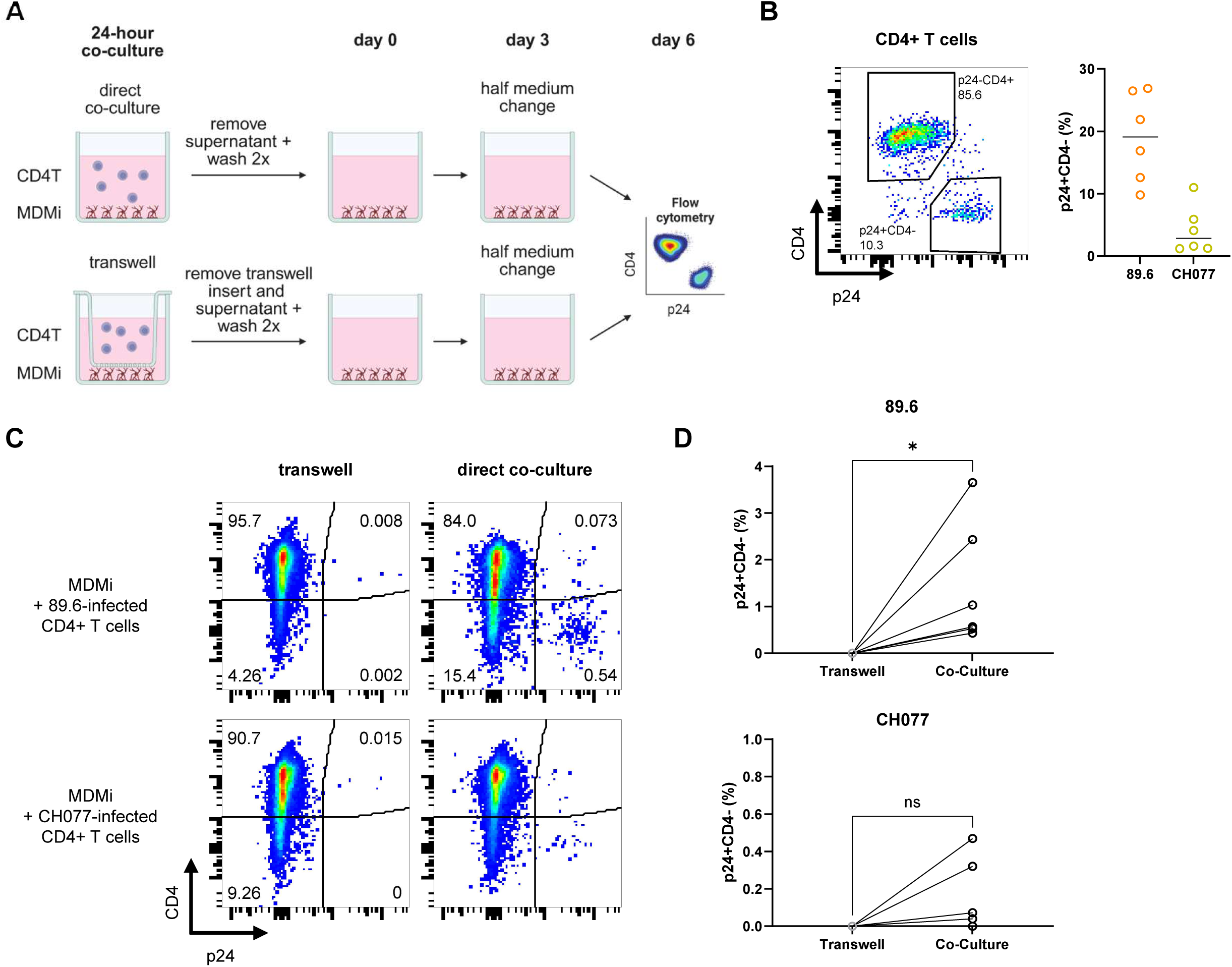
Cell contact between HIV-1-infected CD4+ T cells and MDMi enhances productive MDMi infection. (A) Schematic figure of transwell assay setup: Co-culture of MDMi (red) and HIV-1-infected CD4+ T cells (grey), with or without a transwell insert permitting virion but not immune cell migration, stopped after 24 hours by removal of transwell insert and culture medium. After two washing steps, MDMi culture was continued for 6 days before further analysis. (B) Representative flow cytometry plot and cumulative data of infection frequency of 89.6– and CH077-infected CD4+ T cells before start of the co-culture as determined by the percentage of p24+CD4-cells among CD3+ live single cells. (C) Representative flow cytometry plot and (D) cumulative data of infection frequency of MDMi 6 days post co-culture with 89.6 or CH077-infected CD4+ T cells with or without transwell insert as determined by the percentage of p24+CD4-cells among CD11b+ live single cells. Statistical significance was assessed using the Wilcoxon matched-pairs signed-rank test (ns – not significant; * – p < 0.05).

In the transwell co-culture set-up, permitting only cell-free virus transmission, no MDMi infection was detected with HIV-1 CH077 (n=6) (Figure 2C, D, Supplemental Figure S2A). Likewise, infection with HIV-1 89.6 was undetectable in all but one donor (n=6), in whom an infection frequency of 0.03% was observed. In contrast, co-culture enabling direct cell-to-cell contact was associated with significantly higher infection frequencies for 89.6 (infection frequency 0.43–3.65%; n=6) or detectable infection for 4 out of 6 donors for CH077 (0.04–0.47%). For most donors, higher initial infection frequencies of CD4+ T cells were associated with higher infection frequencies of co-cultured MDMi (Supplemental Figure S2B). To further quantify productive 89.6 infection of MDMi after CD4+ T-cell contact, we assessed p24 nucleocapsid release into the cell culture supernatant after three and six days of microglia culture. p24 release was higher in direct co-culture (2,1197-113,569 pg/mL at day 6) than in the transwell set-up (0-213 pg/mL at day 6) and increased from day 3 to day 6 for most donors (n = 5 out of 6) under co-culture conditions (Supplemental Figure S2C). Altogether, productive HIV-1 infection of MDMi was enhanced through direct cell contact between infected CD4+ T cells and MDMi targets.

### HIV-1 infection modulates CD4+ T-cell ligands that engage key receptors on MDMi

Phagocytosis, heterotypic cell fusion, tunneling nanotubes and virological synapse-like transfer have been proposed as mechanisms of cell contact-dependent HIV-1 transmission from CD4+ T cells to macrophages. Having observed enhanced infection of MDMi following direct contact with CD4+ T cells, we next aimed to assess these potential mechanisms by analyzing gene expression and HIV-1-induced surface protein changes of relevant ligand/receptor axes in microglia–CD4+ T cell crosstalk.

Within CD4+ T cell cultures exposed to either HIV-1 strain 89.6 or CH077, the productively infected p24+CD4-population expressed significantly lower surface levels of the “don’t-eat-me” molecule CD47 compared to the uninfected p24−CD4+ population (Figure 3A). To next determine whether CD4+ T cells exposed to HIV-1 89.6 or CH077 also acquire an “eat me” phenotype, phosphatidylserine (PtdSer) exposure was assessed by Annexin V staining. 7-Amino-Actinomycin D (7-AAD) staining was used to exclude irreversibly apoptotic and non-viable cells, with CD4+ T cells incubated at 65°C serving as a positive control. Exposure to 89.6 was associated with a significantly increased frequency of Annexin V+7-AAD− cells, whereas exposure to CH077 did not significantly alter the frequency of this population (Figure 3B). Subsequent intracellular p24 staining of sorted 89.6-exposed CD4+ T cells showed enrichment of infected p24+ CD4+ T cells (36.6–51.2%) in the Annexin V+7-AAD− population (Supplemental Figure S3A-D). We next confirmed that the MDMi model expressed both CD172a (SIRPα), the cognate receptor for CD47, and the PtdSer receptors TREM-2 and MerTK at the cell surface (Figure 3C).

**Figure 3:**
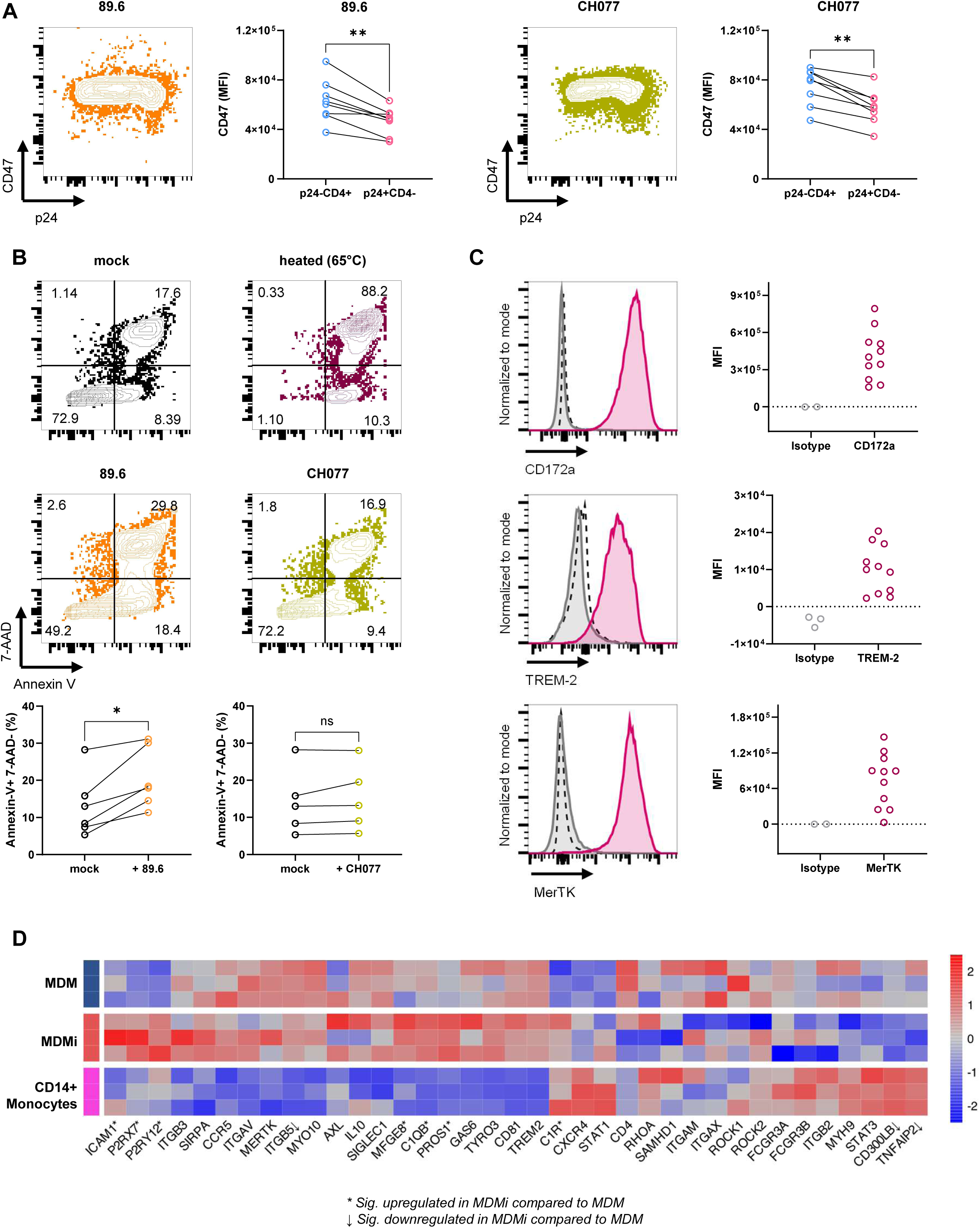
HIV-1 infection modulates CD4+ T-cell ligands that engage key receptors on MDMi. (A) Representative flow cytometry plots of surface CD47 expression and intracellular p24 staining 3 days post infection of primary CD4+ T cells with HIV-1 89.6 or CH077 and cumulative data of CD47 surface expression (MFI). Statistical significance was assessed using the Wilcoxon matched-pairs signed-rank test (** – p < 0.01). (B) Representative flow cytometry plots of Annexin V and 7-AAD staining 3 days post infection of primary CD4+ T cells with HIV-1 89.6 or CH077 and cumulative data of percentage of Annexin V+7-AAD– among single cells. Heated cells (25 minutes at 65 °C) served as positive control for apoptosis. Statistical significance was assessed using the Wilcoxon matched-pairs signed-rank test (ns – not significant; * – p < 0.05). (C) Representative histograms (dotted line = unstained control, grey = isotype control, red = stained sample) and cumulative data (MFI) of flow cytometric staining of surface CD172a, TREM2 and MerTK expression compared to respective isotype control. (D) Heat map of row-scaled, variance-stabilized bulk RNA-seq expression for selected immune interaction and phagocytosis genes in donor-matched CD14+ monocytes, M-CSF-derived macrophages (MDM) and MDMi from three donors. MDMi and MDM shared a broad myeloid phagocytic signature, whereas MDMi showed higher relative expression of *C1QB, C1R, MFGE8, PROS1, P2RY12, P2RX7* and *ICAM1,* and lower expression of *CD300LB, ITGB5 and TNFAIP2.* Colors indicate relative expression; * and ↓ denote genes significantly upregulated or downregulated, respectively, in MDMi versus MDM.

Bulk RNA-seq further supported a phagocytosis-competent MDMi phenotype that was largely similar to, yet transcriptionally distinct from, M-CSF-derived MDM. Both MDMi and MDM expressed a broad myeloid phagocytic program (such as *ITGAM*, *ITGB2*, *ITGAX*, *TREM2*, *SIRPA*, and *SIGLEC1*), but MDMi were distinguished from MDM by higher expression of genes associated with PtdSer sensing, complement recognition, and microglia-associated purinergic signaling, including *C1QB, C1R, P2RY12,* and *P2RX7* (Figure 3D). MDMi also expressed transcripts involved in PtdSer-dependent efferocytosis, including *MERTK, AXL, TYRO3, GAS6, PROS1, MFGE8, ITGAV, ITGB3,* and *ITGB5*, whereas *CD300LB* was significantly reduced relative to M-CSF-derived MDM. In addition, MDMi expressed candidate genes associated with adhesion, virus capture, and cytoskeletal interactions, including *ICAM1, SIGLEC1, MYO10,* and *RHOG*, that may potentially be involved in contact-dependent routes of HIV-1 transfer beyond cell-free infection.

Taken together, these data indicate that HIV-1 infection shifts CD4+ T cells toward an engulfment-prone phenotype characterized by reduced CD47 expression and increased PtdSer exposure, while MDMi display a distinct phagocytic phenotype that may enable recognition and uptake of infected T cells.

### HIV-1 infection of CD4+ T cells promotes cell–cell contact with MDMi

Having observed HIV-1-induced modulation of potential ligand/receptor pairs between MDMi and infected CD4+ T cells, we next aimed to investigate and visualize the crosstalk between MDMi and CD4+ T cells. To this end, MDMi and autologous HIV-1-infected CD4+ T cells (infection frequencies 20.6–26%, Supplemental Figure S4A) were co-cultured for one hour to assess interactions via imaging and flow cytometry. Fluorescence microscopy and IncuCyte imaging of fixed and stained cells showed interactions, that remained stable throughout multiple washing steps, between activated and 89.6-infected CD4+ T cells with MDMi (Figure 4A). These interactions were quantified using IncuCyte image analysis of 49 images per condition and donor (Figure 4B). Notably, primary unstimulated CD4+ T cells and MDMi interacted only marginally. Prior activation of CD4+ T cells increased the co-localization of MDMi and T cells, and HIV-1 89.6 infection further enhanced cell–cell interactions of MDMi with both p24+ HIV-1-infected CD4+ T cells and uninfected bystander cells. We next assessed whether these interactions led to increased engulfment of CD4+ T cells by MDMi. To this end, we established a flow cytometry-based gating strategy to quantify T cell uptake by MDM by staining CD11b+ MDM with extracellular CD3-AF488 (before permeabilization), to identify T cells attached on the outside, and intracellular CD3-AF647 (after permeabilization), enabling detection of all T cells including T cells engulfed by MDM (Supplemental Figure S4B) [51]. As a positive control for phagocytosis, antibody-coated CD4+ T cells inducing antibody-dependent cellular phagocytosis (ADCP) were used, resulting in almost 50% of MDM being single-positive for intracellular CD3. This effect was strongly reduced at 4 °C – a temperature that permits binding of T cells but prevents their uptake by MDM. Activation of T cells substantially increased CD3 signal in MDM (double positive for extracellular and intracellular CD3) suggesting enhanced interaction of T cells with MDM. In contrast, the fraction of MDM positive only for intracellular CD3, potentially reflecting engulfment, was not markedly changed compared to unstimulated CD4 T cells. This internalization was also drastically reduced in the 4 °C negative control (Supplemental Figure S4B). Using the same gating strategy, MDMi were first identified by CD11b staining and subsequently gated on extra– and intracellular CD3 staining (Supplemental Figure S4C). In line with the IncuCyte data, activated CD4+ T cells showed increased attachment to MDMi compared to unstimulated T cells (CD3intr+ CD3extr+ CD11b+ cells) (Figure 4C, D). However, co-culture with 89.6-infected CD4+ T cells increased the proportion of MDMi positive only for intracellular CD3, as a proxy for uptake. Therefore, we next used ImageStream technology that combines flow cytometry with high-resolution imaging of single cell interactions to better understand the spatial localization of interacting CD4+ T cells with MDMi observed in our flow cytometry-based assay. ImageStream analysis revealed diverse interactions between co-cultured MDMi and 89.6-infected CD4+ T cells, including (i) the co-localization of CD3 and p24 at the MDMi surface, indicating the beginning of a heterotypic cell fusion process, (ii) a lack of CD3 surface staining at the MDMi-CD4+ T cell interface, suggesting virological synapse formation with limited antibody accessibility or an early fusion or phagocytosis event, and (iii) MDMi surface membrane extending toward a T cell, potentially displaying early cell fusion or ongoing phagocytic uptake (Figure 4E). We next proceeded to live-cell imaging to capture the MDMi–CD4+ T cell interactions dynamically using infection with the transmitted founder virus CH077. Live-cell imaging of MDMi co-cultured with CH077-infected and bystander CD4+ T cells showed a T cell and MDMi getting in contact, followed by T cell death and subsequent phagocytosis by the MDMi (Figure 4F). Altogether, these data demonstrate that activation and HIV-1 infection of primary CD4+ T cells result in increased interaction with MDMi, including engulfment of HIV-1-infected CD4+ T cells.

**Figure 4:**
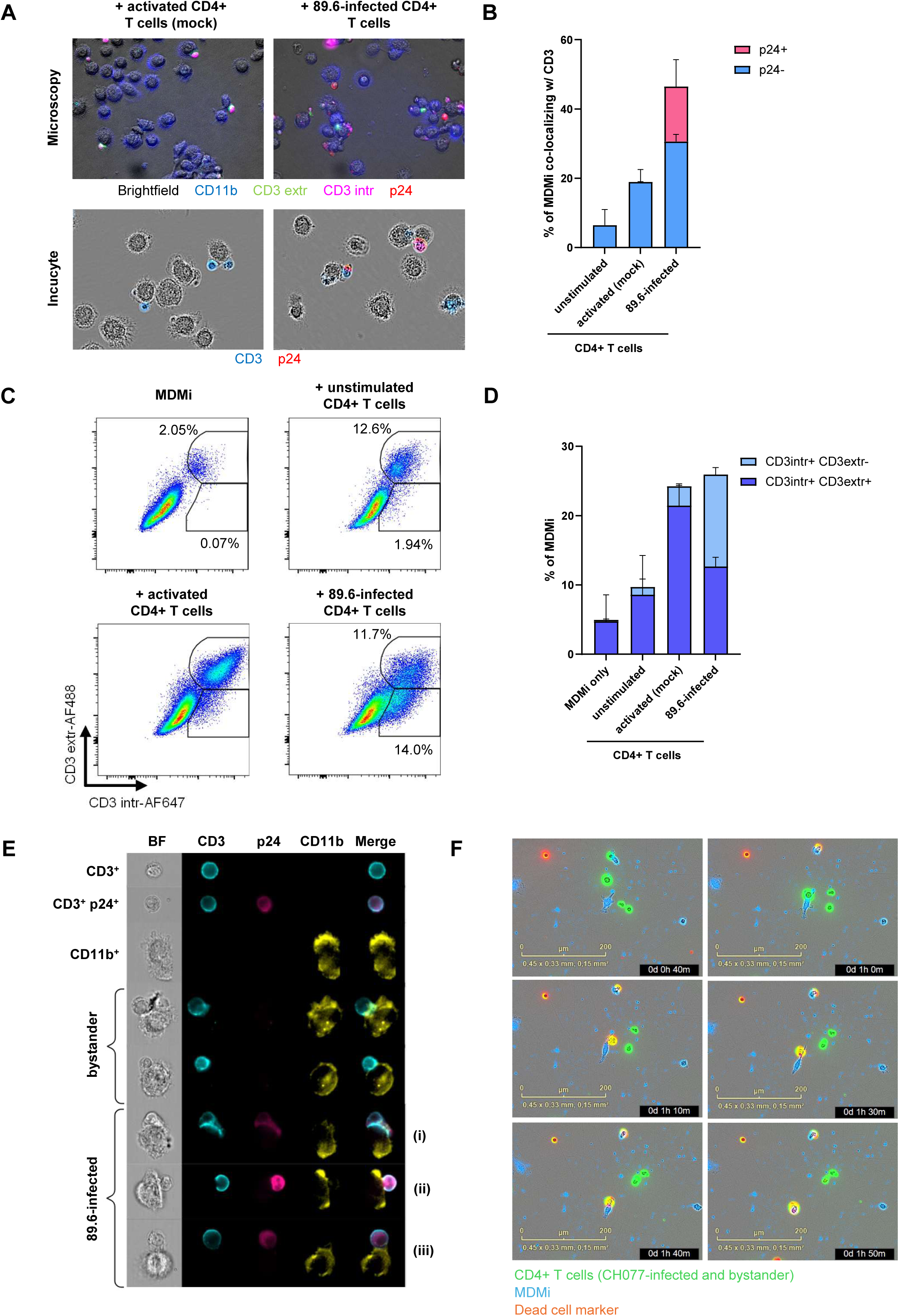
HIV-1 infection of CD4+ T cells promotes cell-cell contact with MDMi. MDMi were co-cultured for 1 h with unstimulated, activated (mock) and 89.6-infected autologous CD4+ T cells, then washed, stained and analyzed via microscopy, IncuCyte, flow cytometry and ImageStream. (A) Representative immunofluorescence images showing merged brightfield, CD11b, extracellular CD3, intracellular CD3 and HIV p24 channels (upper row) and representative IncuCyte images including CD3 and p24 markers (lower row). (B) Quantitative IncuCyte image analysis of MDMi (selected by size) co-localizing with CD3 signal, divided into p24+ and p24− objects (49 images per condition and donor). Bars represent mean + standard deviation of n = 2 donors. (C) Representative flow cytometric plots showing extracellular CD3-AF488 and intracellular CD3-AF647 staining of MDMi and MDMi–T cell conjugates after gating on CD11b+ cells. (D) Quantification of flow cytometric analysis of MDMi and conjugates that are intracellular CD3+ extracellular CD3– (light blue) and intracellular CD3+ extracellular CD3+ (dark blue). Bars represent mean + standard deviation of n = 2 donors. (E) Representative ImageStream images of uninfected and p24+ T cells, MDMi alone and MDMi in contact with 89.6-infected and bystander T cells. Brightfield, fluorescence, and merged images are shown. Live-cell imaging of MDMi co-cultured with CH077-infected and bystander CD4+ T cells is shown in (F) as representative time-lapse images from IncuCyte live-cell imaging (MDMi in blue, T cells in green, cell death indicator in red).

## Discussion

Our co-culture studies of primary monocyte-derived microglia and autologous, *in vitro* HIV-1-infected CD4+ T cells revealed three key findings: First, while MDMi are permissive to cell-free virus infection, direct cell–cell contact markedly enhanced productive HIV-1 infection of MDMi. Second, both MDMi and HIV-1-infected CD4+ T cells displayed a pro-interaction, pro-phagocytic phenotype. Third, HIV-1 infection increased interaction between CD4+ T cells and MDMi, supporting a model in which contact-dependent mechanisms facilitate viral transfer from CD4+ T cells to microglia.

As sentinels of CNS immune surveillance, microglia restrict cerebral infection through multiple effector mechanisms, including phagocytic clearance of pathogen-infected cells [52]. Consistently, the microglia model expressed receptors for “eat-me” and “don’t-eat-me” signals, such as PtdSer receptors and CD172a, at both transcript and surface protein levels. Multiple viruses, including HIV-1 and influenza A virus, are known to hijack phagocytic pathways to facilitate infection of myeloid cells [18, 53, 54]. Here, HIV-1 infection of primary CD4+ T cells with both the cell-culture adapted strain 89.6 and the transmitted/founder virus CH077 downmodulated surface levels of CD47, the cognate ligand of CD172a, concordant with prior reports [19]. Modulation of the CD47-CD172a axis may therefore contribute to HIV-1 cell-to-cell infection. In a murine model, increased PtdSer exposure on cells transduced with adenovirus type 5-based vectors promoted their recognition and phagocytosis by microglia [55]. In line with prior data describing PtdSer exposure on CD4+ T cells following infection with the cell-culture adapted HIV-1 strains NL4-3 and HXB2 [17, 56], we observed increased PtdSer exposure on live 89.6-infected CD4+ T cells. Phagocytosis of viable HIV-1-infected cells by macrophages has been demonstrated before, however, it was less efficient than phagocytosis of apoptotic infected cells [21, 56]. Within this framework, our findings indicate that HIV-1 infection skews CD4+ T cells toward an engulfment-prone phenotype, potentially contributing to the observed increased interactions with MDMi. The extent to which the CD47-CD172a axis and PtdSer-dependent recognition pathways mediate productive microglial infection and how HIV-1 has evolved to potentially leverage these pathways remains to be fully delineated.

Phagocytosis, heterotypic cell fusion, tunneling nanotubes, and virological synapse-like transfer have been proposed as non-mutually exclusive mechanisms of HIV-1 transmission from CD4+ T cells to myeloid cells [29]. Here, we observed enhanced CD4+ T cell–microglia contact following HIV-1 infection of CD4+ T cells. This is consistent with a previous study demonstrating that HIV-1 transmission from macrophages to CD4+ T cells is accompanied by stabilization and prolongation of macrophage–CD4+ T cell interactions, which was mediated by gp120/CD4 and LFA-1/ICAM-1 engagement [57]. LFA-1 is also needed for CD4+ T cell diapedesis through the blood brain barrier [58] and, consequently, *LFA-1* expression has also been reported in CD4+ T cells disseminating SIV to the CNS [31]. Our data favours a model where activation of CD4+ T cells increases the attachment to MDMi, possibly representing surveillance checks, however, following HIV-1-infection, more of these interactions lead to phagocytosis of infected CD4+ T cells. Studies with clinical samples and SIV models support the concept that tissue macrophages acquire infection through engulfment of infected CD4+ T cells by detecting T cell receptor DNA in both alveolar macrophages from people living with HIV and SIV+ myeloid cells in the rhesus macaque model [14, 15]. Furthermore, phagocytosis of HIV-1-infected CD4+ T cells by MDMs and alveolar macrophages resulting in productive infection has been demonstrated conclusively *in vitro* [16, 21]. The increased cell–cell contact between MDMi and both HIV-1-infected and bystander CD4+ T cells, observed across multiple imaging modalities, likely reflects a range of interaction mechanisms, including early phagocytosis, heterotypic cell fusion, and virological synapse-like contacts. In clinical histopathology, multinucleated giant cells are regarded as a hallmark of HIV encephalitis and are closely linked with HIV-associated dementia [2, 59]. Mascarau et al. (2023) showed that phagocytosis-mediated infection of monocyte-derived macrophages (MDMs) predominantly occurs when HIV-1-infected T cells are apoptotic, whereas infection via heterotypic cell fusion with viable infected CD4+ T cells is more efficient and results in higher infection frequencies [25]. The same study further provided evidence for heterotypic cell fusion across multiple tissue macrophage populations but did not investigate the brain. Notably, 89.6 – the HIV-1 variant used in our co-culture experiments – has been characterized as highly cytopathic [45] and is known to induce syncytium formation [60], potentially facilitating both phagocytosis and heterotypic cell fusion. Heterotypic cell fusion involves gp120/CD4 interactions, theoretically restricting infection of myeloid cells through this mechanism to M-tropic HIV-1 strains adapted to low CD4 surface density [23]. However, fusion of MDM with infected CD4+ T cells has been reported to enhance the efficiency of CD4 and CCR5 usage, thereby enabling infection of MDM by T-tropic HIV-1 strains [24]. In contrast to phagocytosis, cell fusion is susceptible to blocking via a subset of neutralizing antibodies [61]. Therefore, further studies are needed to dissect the relative contributions of these individual mechanisms to HIV-1 spread from CD4+ T cells to microglia.

Microglia have now been conclusively demonstrated to harbour the replication competent viral reservoir in the brain, with published data from post-mortem brain tissue of a virologically suppressed individual, which reported an intact proviral reservoir of 1,111-1,985 copies per million microglia [62]. In the same study, 80% of isolated virus clones replicated efficiently in both microglia and CD4+ T cells [62]. Our study provides a proof-of-concept that HIV-1 can be transmitted *in vitro* from CD4+ T cells to microglia-like cells, leading to productive infection. Inoculation with cell-free virus stocks yielded productive infection of microglia, however, in the co-culture system, direct cell-to-cell contact with infected CD4+ T cells significantly increased microglia infection. Microglial infection during early HIV-1 viremia is unlikely to arise from cell-free virus, given that T-tropic variants requiring high CD4 density dominate during reservoir establishment, whereas myeloid cells express only low surface CD4 [11, 12]. In line with our data, co-culture studies using alveolar macrophages have similarly reported increased p24 release in culture supernatants, consistent with enhanced productive infection through direct cell–cell contact [16]. Notably, addition of uninfected CD4+ T cells to macrophage cultures infected with patient-derived HIV-1 enhanced viral replication and was associated with increased formation of virus-containing compartments [63]. The similarity of HIV-1 sequences between the brain and peripheral blood [11, 33, 64], along with the presence of CCR5+ CD4+ T cells throughout the brain and a distinct SIV-enriched myeloid/T cell transcriptional cluster identified in the CNS of SIV-infected rhesus macaques [31, 32], together with our findings, are consistent with a role for HIV-1 transmission from CD4+ T cells to microglia in the brain.

Because HIV-1 neuroinvasion occurs early during acute infection, preventing CNS entry by blocking migration of infected CD4+ T cells from the peripheral blood into the brain is unlikely to be clinically feasible. Accordingly, prevention of neurological disease is likely to depend primarily on limiting viral replication and spread within the brain. However, cell-to-cell spread is not fully targeted by antiretroviral therapy, and HIV-1 transmission from CD4+ T cells to myeloid cells via heterotypic cell fusion can evade type I interferon responses and host restriction factors [23, 37, 38]. Other therapeutic approaches, such as broadly neutralizing antibodies targeting the HIV-1 envelope glycoprotein on the surface of infected cells, have shown reduced efficacy against cell-to-cell transmission compared with cell-free infection, possibly reflecting differences in Env accessibility [65]. In this study, we demonstrate *in vitro* HIV-1 transmission from CD4+ T cells to microglia-like cells and show that HIV-1 infection of CD4+ T cells enhances CD4+ T cell–microglia interactions. Additionally, we comprehensively characterized the monocyte-derived microglia model as an accessible resource for studies of the human myeloid HIV-1 brain reservoir. Further mechanistic studies of these interactions in tissue and in particular in the brain will be required to identify potential inhibitors of cell-to-cell transmission of HIV-1.

## Acknowledgements

We thank the laboratories of Beatrice Hahn, John C. Kappes, Frank Kirchhoff and Daniel Sauter for generating and providing the proviral HIV-1 infectious molecular clones Thro.c-GFP and CH077. The following reagent was obtained through the NIH HIV Reagent Program, Division of AIDS, NIAID, NIH: Human Immunodeficiency Virus Type 1 89.6 Infectious Molecular Clone (p89.6), ARP-3552, contributed by Dr. Ronald G. Collman. The following reagent was obtained through the NIH HIV Reagent Program, Division of AIDS, NIAID, NIH: TZM-bl cell line (#8129), contributed by Dr. David Montefiori and Dr. Gabriel Perez. Furthermore, we thank Arne Düsedau, head of the technology platform Flow Cytometry/FACS, and Patrick Blümke, head of the technology platform Next Generation Sequencing, at the Leibniz Institute of Virology for technical support, as well as the healthy cohort coordinators and donors. A.H., K.S. were supported by funding from the Federal Ministry of Research, Technology and Space (BMFTR, 01KI2110). A.H. received funding from the Deutsche Forschungsgemeinschaft (DFG, German Research Foundation) – SFB 1648/1 2024–512741711. A.H. is associated with the iSTAR program from the Federal Ministry of Research, Technology and Space. F.R. received a clinical-leave grant from the German Center for Infection Research (TI 07.001_009). M.G.L. was supported by the Else Kröner-Fresenius-Stiftung iPRIME Scholarship (2021_EKPK.10), UKE, Hamburg. This study was supported by the 3R (Replace, Reduce, Refine) Start-up Funding Program, awarded by the Medical Faculty Hamburg in 2021 to A.H and M.A.F.. This study was supported by the 3R (Replace, Reduce, Refine) Start-up Funding Program, awarded by the Medical Faculty Hamburg in 2018 to S.K.. K.S. was supported by a research grant for doctoral students by the German Academic Exchange Service (DAAD, 57745342).

## Author Contributions

Conceptualization, A.H., and W.G.-B.; methodology, F.R., K.S., M.G.L., L.M., S.K., M.A.F and A.H.; formal analysis, F.R., K.S., and M.G.L.; investigation, F.R., K.S., M.G.L., M. Adiba, T.V.K., J.B.E., U.L. and S.S.; resources, W.G.-B, M. Altfeld and A.H.; data curation, F.R., M.G.L., and K.S.; writing original draft, F.R., and A.H.; writing – review and editing, F.R., K.S., M.G.L., L.M., M. Adiba, T.V.K., J.B.E., I.L., L.B., S.S., M. Altfeld, M.A.F., S.K., U.C.L., W.G.-B., and A.H.; visualization, F.R., K.S., M.G.L.; supervision, W.G.-B, A.H.; funding acquisition, F.R., K.S., M.G.L. and A.H..

## Figure Legends Supplemental Figures

**Supplemental Figure 1:**
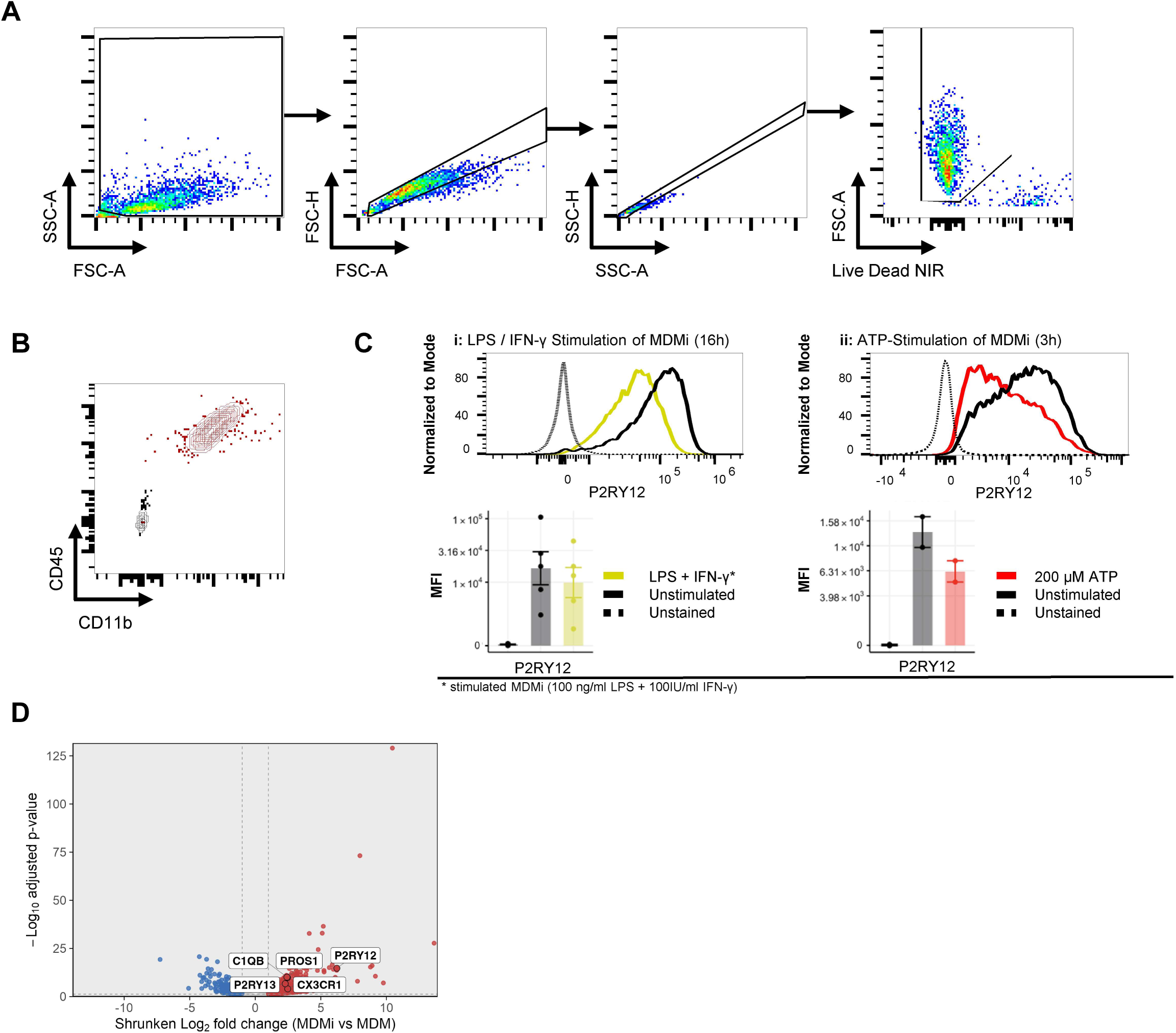
Characterization of monocyte-derived microglia (MDMi) (A) Representative plots demonstrating flow cytometric gating strategy for MDMi: sequential gating on cells, single cells and live cells. (B) Representative flow cytometry plot of CD45 and CD11b surface expression on MDMi (red = stained MDMi, black = unstained controls). (C) Stimulation of MDMi with either 100 ng/mL LPS plus 100 IU/mL IFN-γ for 16 h or 200 µM ATP for 3 h resulted in reduced surface expression of P2RY12 compared with unstimulated cells, consistent with activation-associated downregulation of this microglial marker. Representative flow-cytometry histograms are shown above, with corresponding quantification of P2RY12 median fluorescence intensity (MFI) below. Histograms were normalized to mode, dotted lines indicate unstained controls, and dots represent individual biological replicates. (D) Volcano plot derived from bulk RNASeq analysis (Figure 1A) depicting differential gene expression of MDM and MDMi from 3 donors, highlighting *P2RY12, P2RY13, CX3CR1, C1QB*, and *PROS1* transcripts.

**Supplemental Figure 2:**
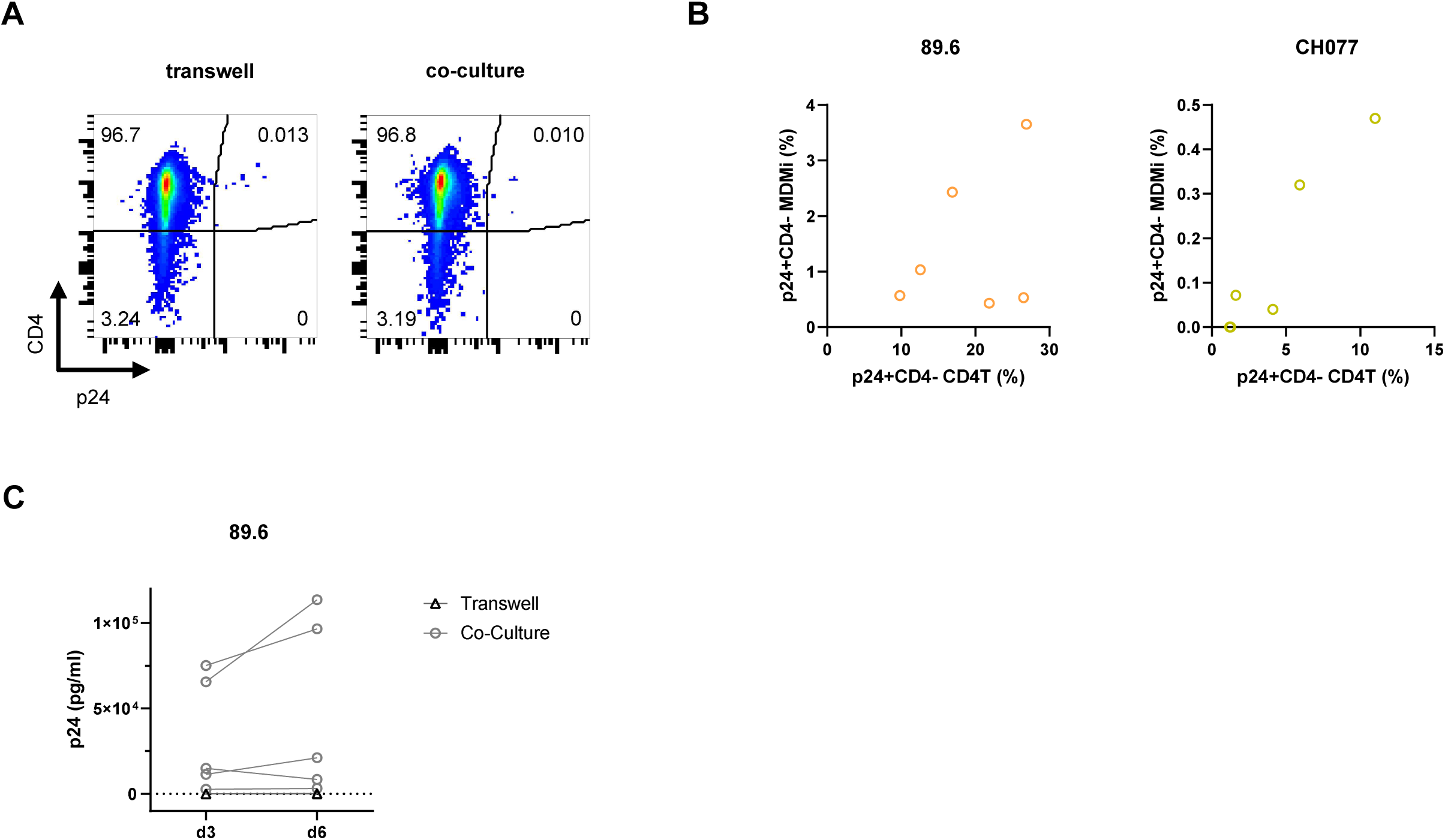
MDMi infected with HIV-1 through co-culture with infected CD4+ T cells release the HIV nucleocapsid p24. (A) Representative flow cytometry plot of MDMi 6 days post co-culture with uninfected CD4+ T cells with or without transwell insert. (B) Comparison of infection frequency (percentage of p24+CD4-cells) between CD4+ T cells (CD4T) prior to co-culture and MDMi 6 days post co-culture. (C) Concentration of HIV nucleocapsid p24 in the supernatant of MDMi cultures 3 and 6 days post co-culture with HIV-1 89.6-infected CD4+ T cells with or without transwell insert.

**Supplemental Figure 3:**
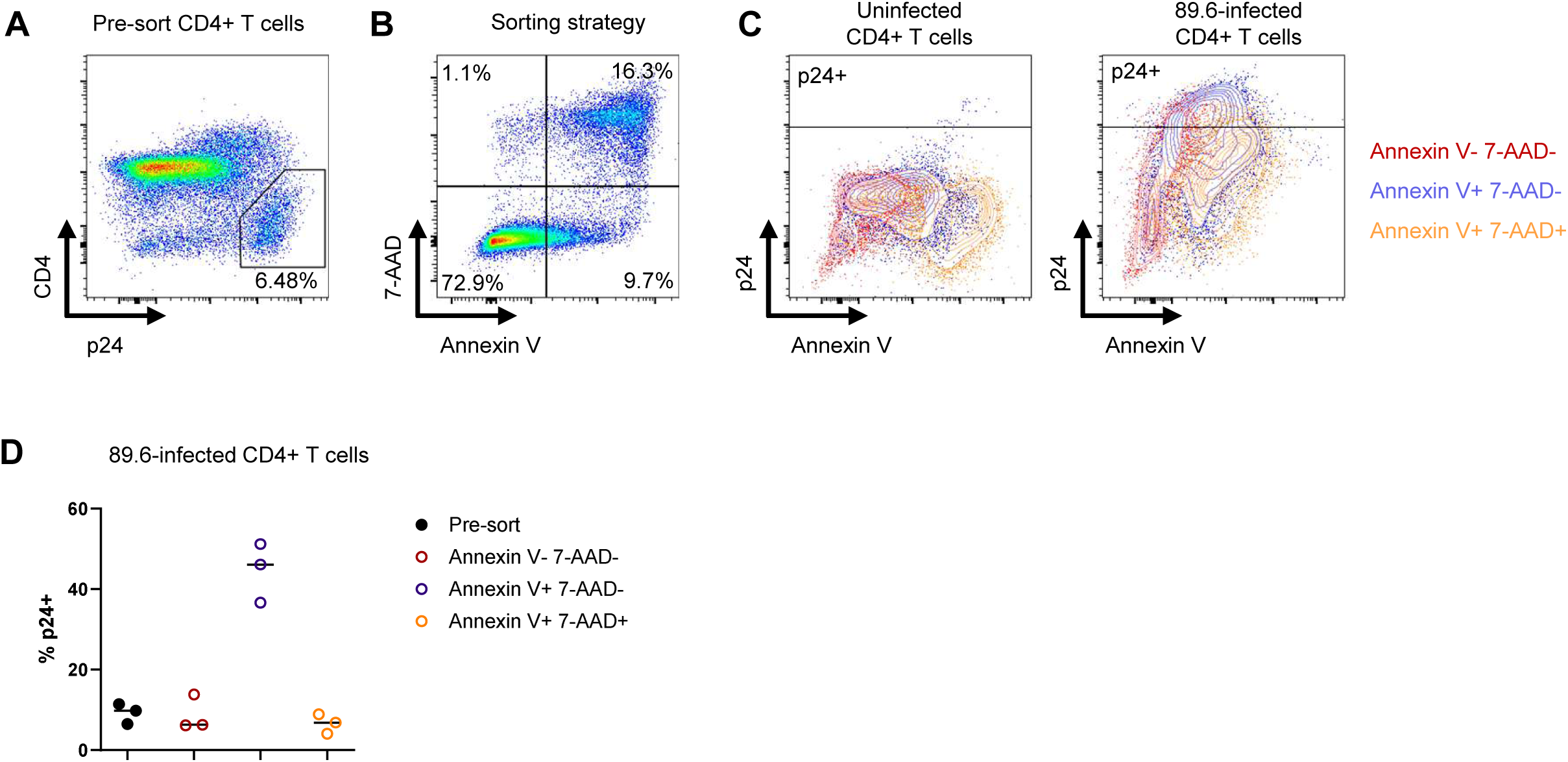
HIV-1-infected p24+ CD4− T cells are enriched in the Annexin V+/7-AAD− population. (A) Representative flow cytometric gating of infected CD4+ T cells pre-sorting. (B) Representative flow cytometric plot showing the sorting strategy based on Annexin V and 7-AAD signals. (C) Representative flow cytometric plots displaying the fluorescence intensity for p24 and Annexin V of the sorted Annexin V−7-AAD−, Annexin V+7-AAD− and Annexin V+7-AAD+ populations, comparing uninfected and 89.6-infected CD4+ T cells. (D) The percentage of p24+ cells within the 89.6-infected T cell population pre-sorting and Annexin V−7-AAD−, Annexin V+7-AAD− and Annexin V+7-AAD+ populations post-sorting as determined by flow cytometry (n = 3 donors).

**Supplemental Figure 4:**
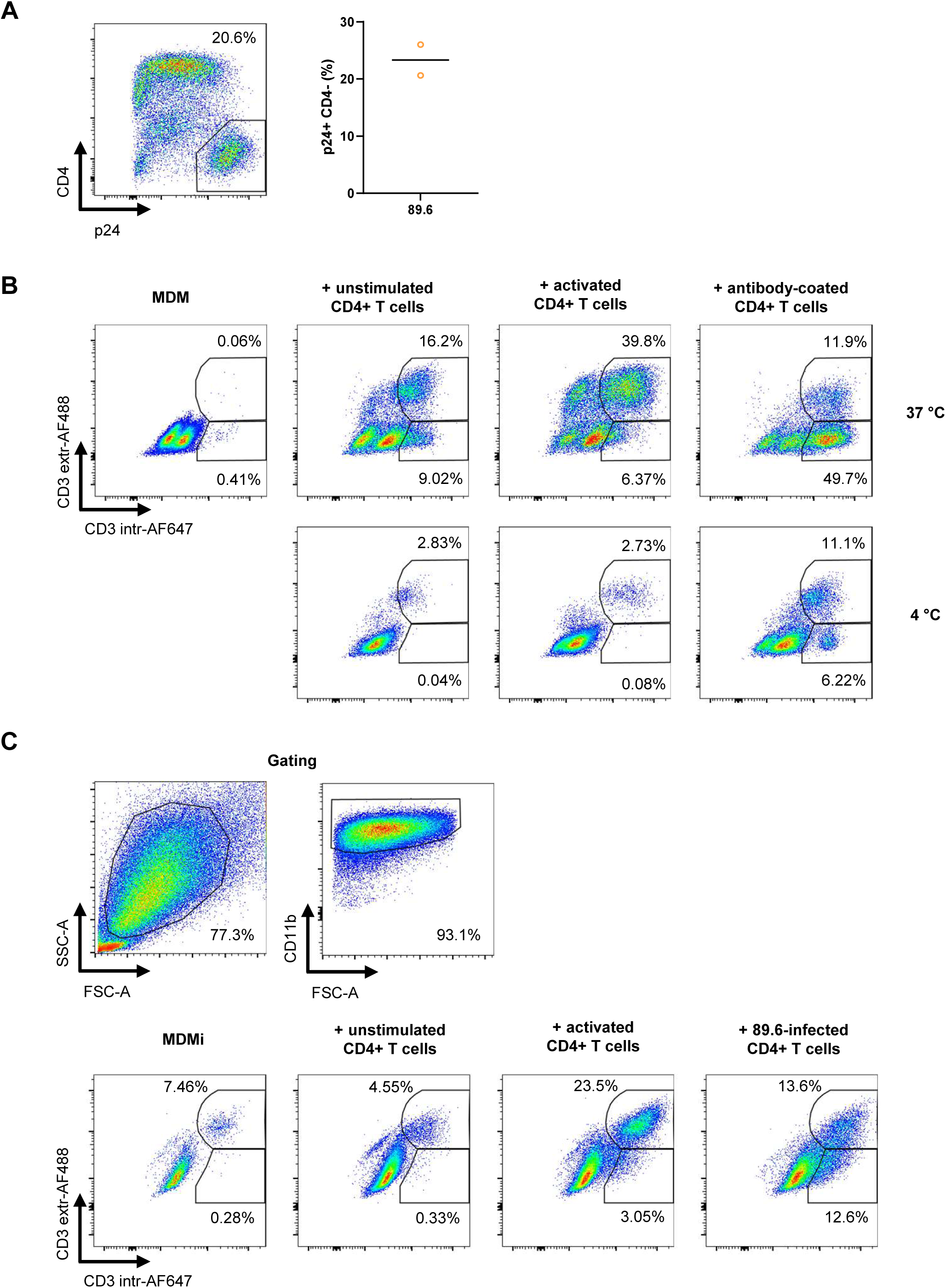
Phagocytosis assay of HIV-1-infect CD4+ T cells with both MDMi and MDM. (A) Gating on HIV-1-infected cells based on p24 positivity and CD4 downregulation and quantification of the infection rates for both donors of the CD4+ T cells 3 days post-infection that were used for MDMi–CD4+ T cell interaction assays (n = 2 donors). (B) Establishment and validation of the extracellular CD3-AF488 and intracellular CD3-AF647 gating strategy using MDM co-cultured for 1 h with autologous CD4+ T cells, including antibody-coated CD4+ T cells (with Ultra-LEAF™ Purified anti-human CD4 Antibody) as a positive control (antibody-dependent phagocytosis) and 4 °C conditions as negative controls. (C) Representative flow cytometric plots showing the gating of MDMi and MDMi–T cell conjugates on FSC/SSC and CD11b positivity and the subsequent extracellular CD3-AF488 and intracellular CD3-AF647 gating after 1 h co-culture of MDMi with unstimulated, activated and 89.6-infected CD4+ T cells for the second donor.

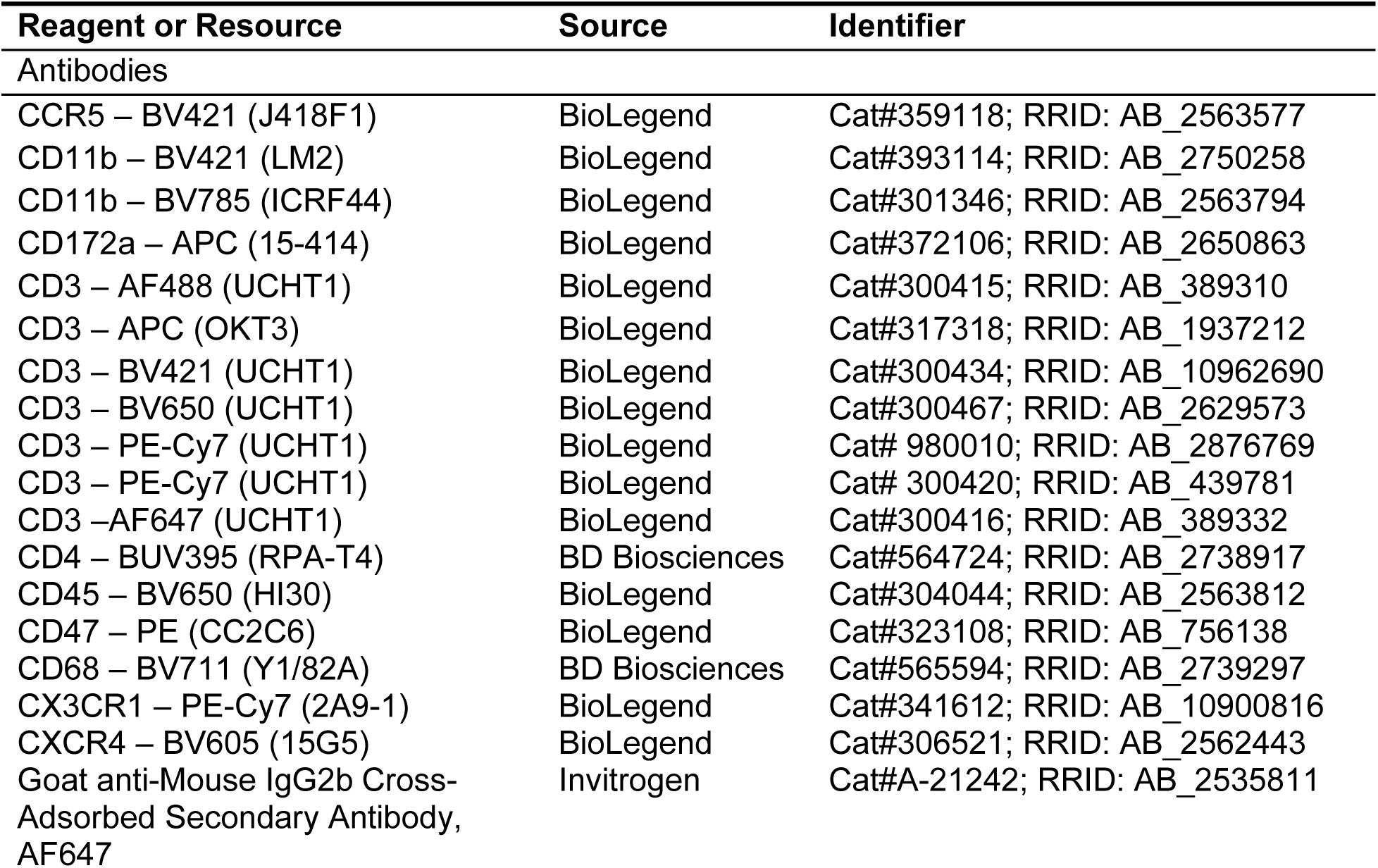
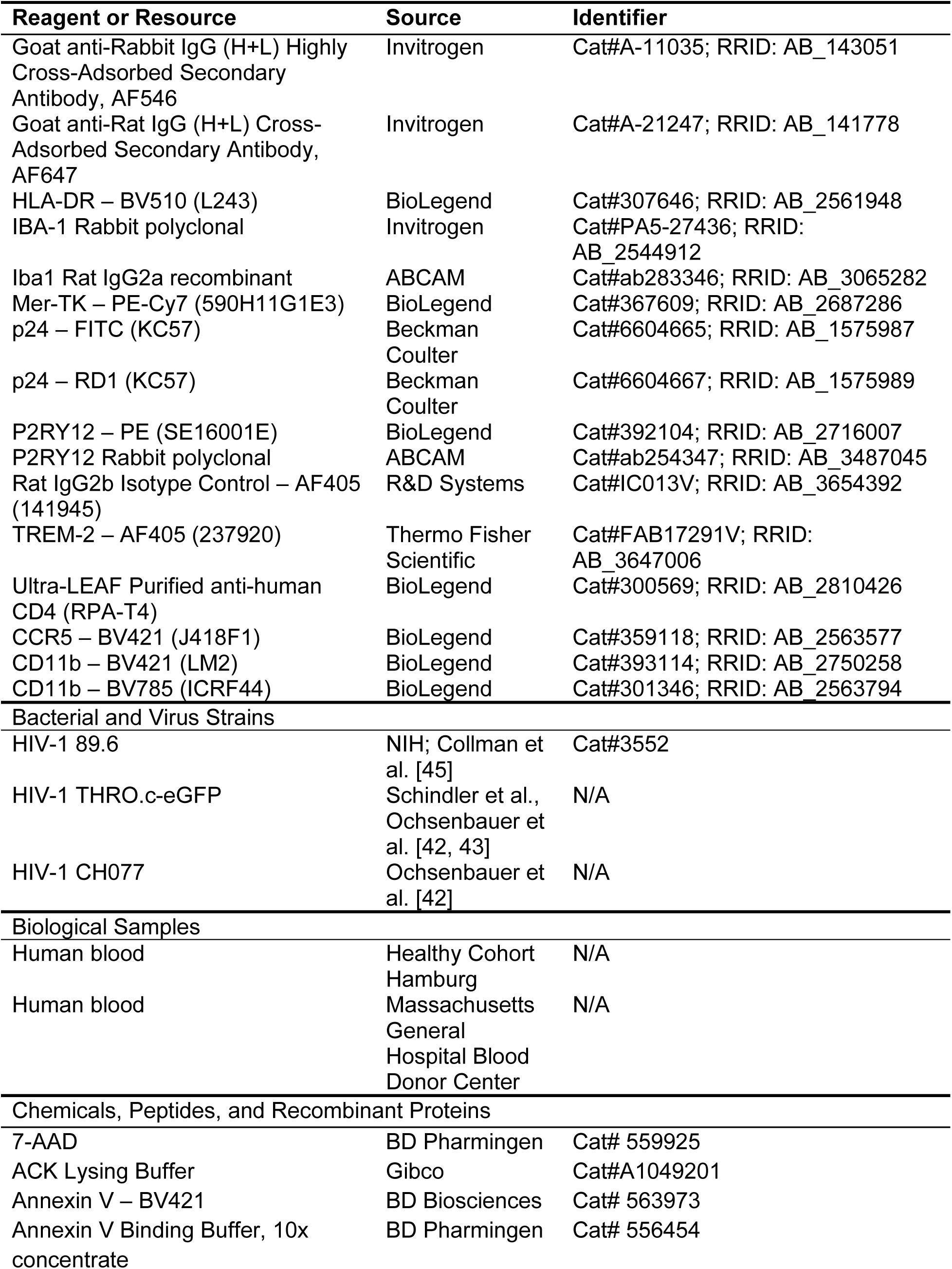
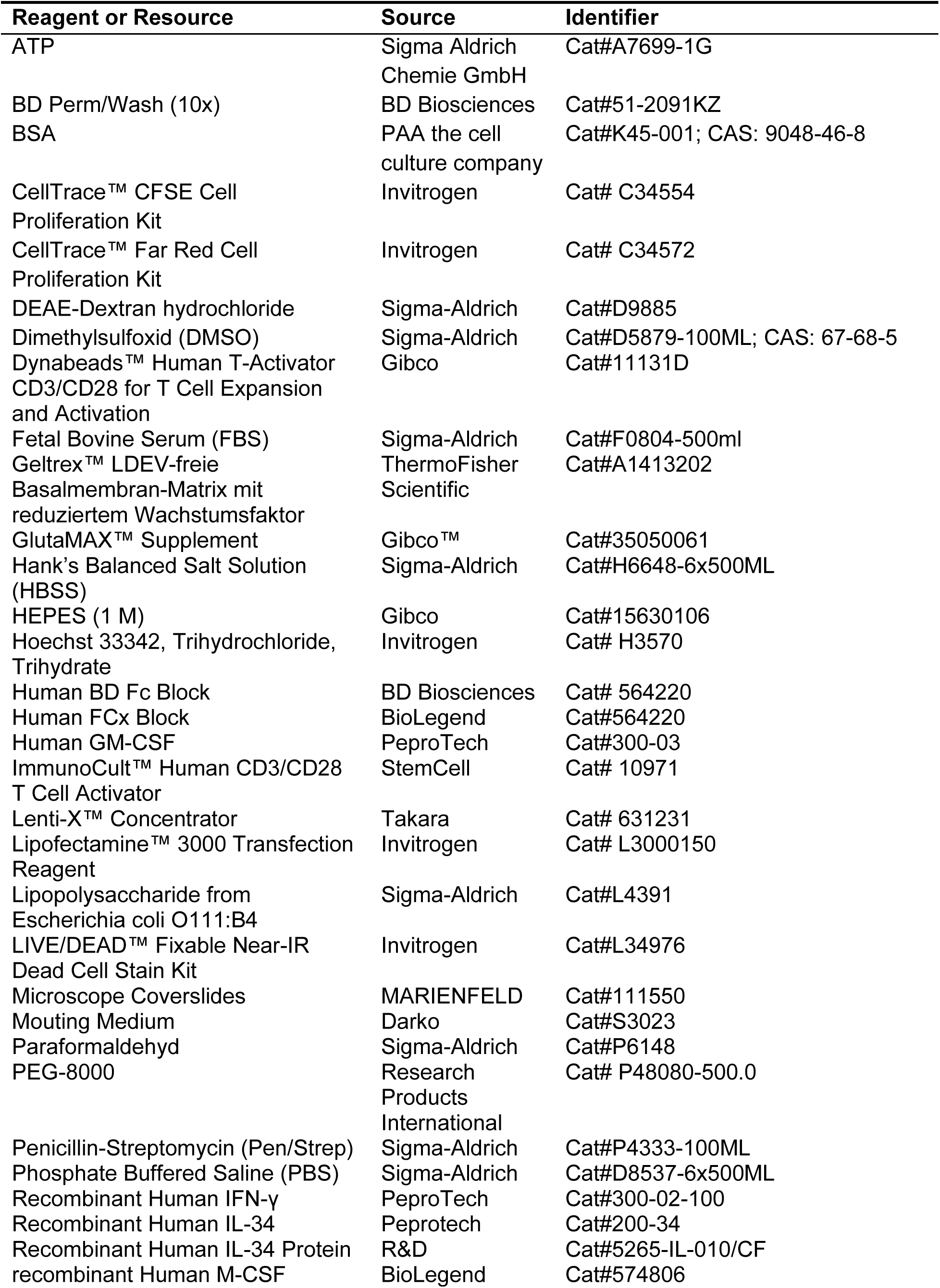
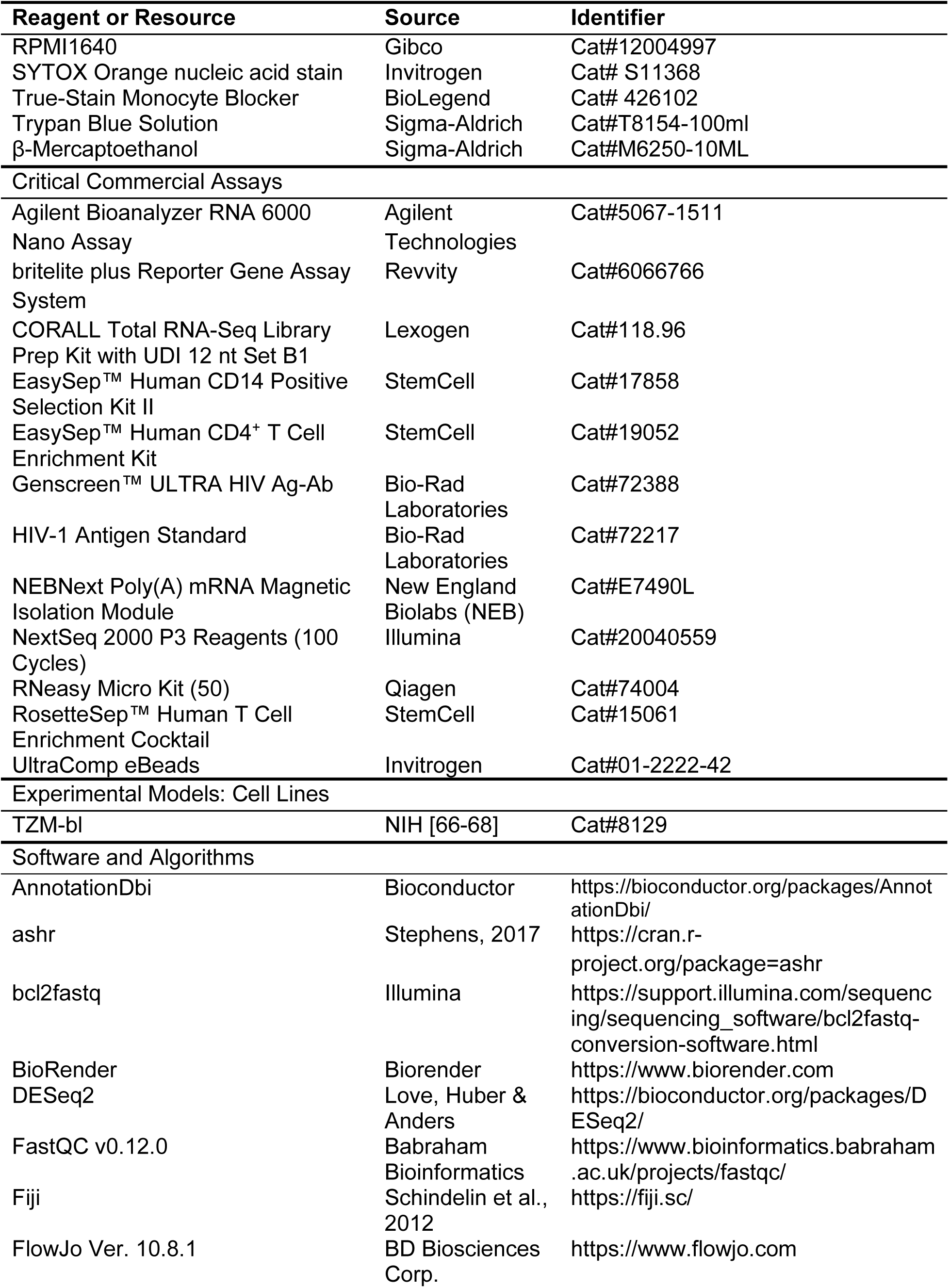
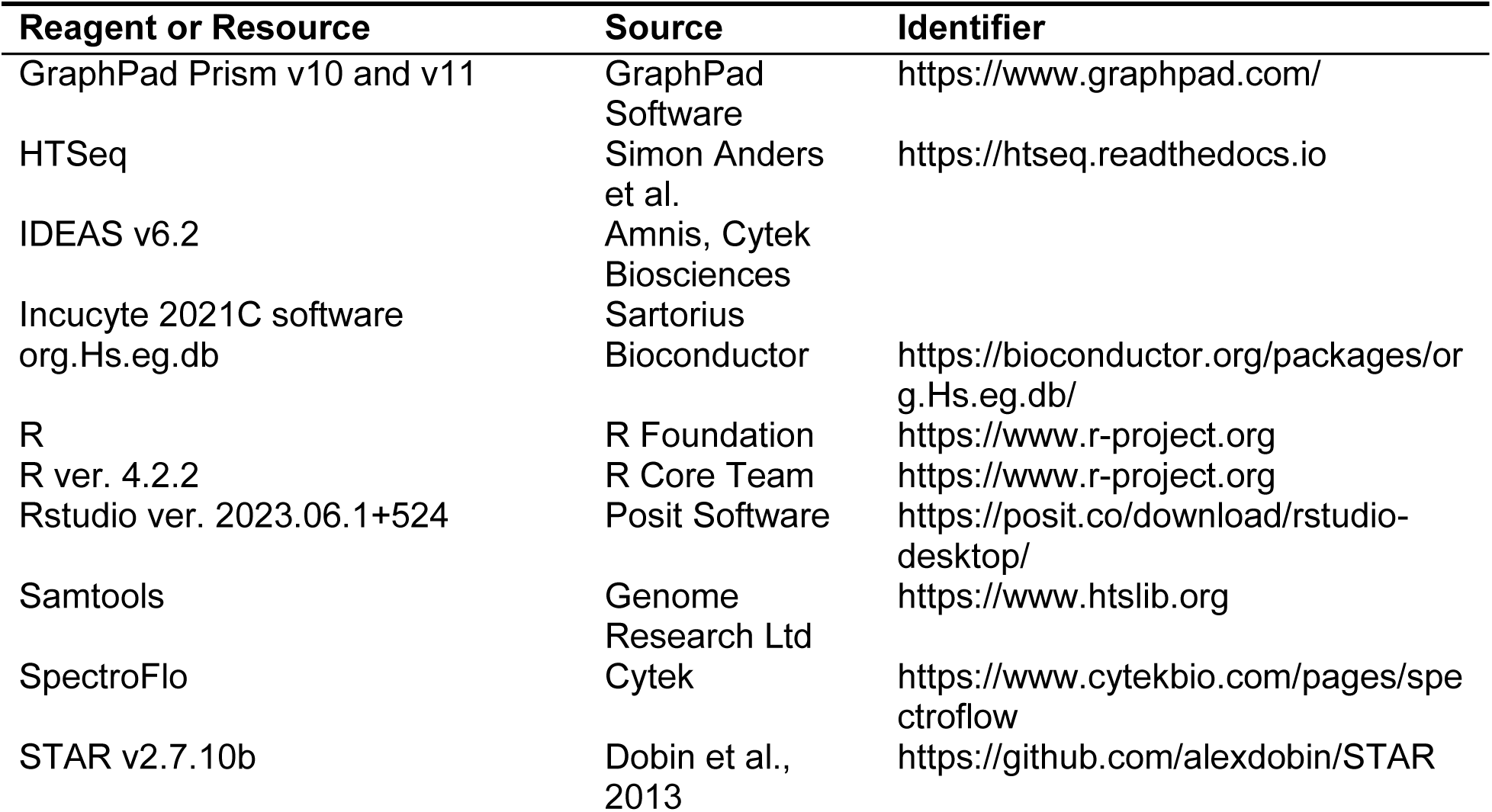
Key Resources Table.

## Figure Legends

